# 3D Printable High Performance Conducting Polymer Hydrogel for All-Hydrogel Bioelectronics

**DOI:** 10.1101/2022.01.29.478311

**Authors:** Tao Zhou, Hyunwoo Yuk, Faqi Hu, Jingjing Wu, Fajuan Tian, Heejung Roh, Zequn Shen, Guoying Gu, Jingkun Xu, Baoyang Lu, Xuanhe Zhao

**Author notes:** These authors contributed equally to this work (Tao Zhou, Hyunwoo Yuk).

## Abstract

Owing to the unique combination of electrical conductivity and tissue-like mechanical properties, conducting polymer hydrogels have emerged as a promising candidate for bioelectronic interfacing with biological systems. However, despite the recent advances, the development of hydrogels with both excellent electrical and mechanical properties in physiological environments remains a lingering challenge. Here, we report a bi-continuous conducting polymer hydrogel (BC-CPH) that simultaneously achieves high electrical conductivity (over 11 S cm^-1^), stretchability (over 400%) and fracture toughness (over 3,300 J m^-2^) in physiological environments, and is readily applicable to advanced fabrication methods including 3D printing. Enabled by the BC-CPH, we further demonstrate multi-material 3D printing of monolithic all-hydrogel bioelectronic interfaces for long-term electrophysiological recording and stimulation of various organs. This study may offer promising materials and a platform for future bioelectronic interfacing.

## Main Text

Electrically conductive hydrogels have emerged as promising alternatives to conventional metallic electrodes in bioelectronics owing to their unique combination of similarity to biological tissues (high water contents, softness) and electrical conductivity^1^. In particular, conducting polymer hydrogels – electrically conductive hydrogels based on conducting polymers – exhibit a set of advantages over other conductive hydrogels based on concentrated ionic salt^2,3^, metal (e.g., Ag, Au, Pt)^4,5^ or carbon nanomaterials (e.g., carbon nanotubes, graphene) ^6^, including favorable electrical properties, stability in physiological environments, biocompatibility, and fully-organic characteristics^1,7,8^.

Despite the recent advances in mechanically robust tough hydrogels mimicking biological tissues^9–13^, the development of mechanically robust conducting polymer hydrogels has faced lingering challenges. Existing tough conducting polymer hydrogels, often prepared by mixing or polymerizing conducting polymers within tough hydrogel matrices, only give low electrical conductivity below 0.3 S cm^-1^ (**Fig. S1**) due to low connectivity between electrical phases in the hydrogels^14–16^. However, attempts to achieve high electrical conductivity by increasing the conducting polymer contents (e.g., pure conducting polymer hydrogels) substantially compromise mechanical properties of hydrogels^17–19^ (**Fig. S1**), limiting their utility in bioelectronic applications that necessitate favorable mechanical and electrical properties simultaneously.

Here we report a bi-continuous conducting polymer hydrogel (BC-CPH) to overcome this challenge. Based on the bi-continuous electrical phase (PEDOT:PSS) and mechanical phase (hydrophilic polyurethane) (**Fig. 1a** and **Fig. S2**), the BC-CPH simultaneously achieves high electrical conductivity (over 11 S cm^-1^), stretchability (over 400%), fracture toughness (over 3,300 J m^-2^), water contents (~ 80%), and tissue-like softness (Young’s modulus below 1 MPa) in physiological environments (**Figs. S1** and **S3**).

**Fig. 1.**
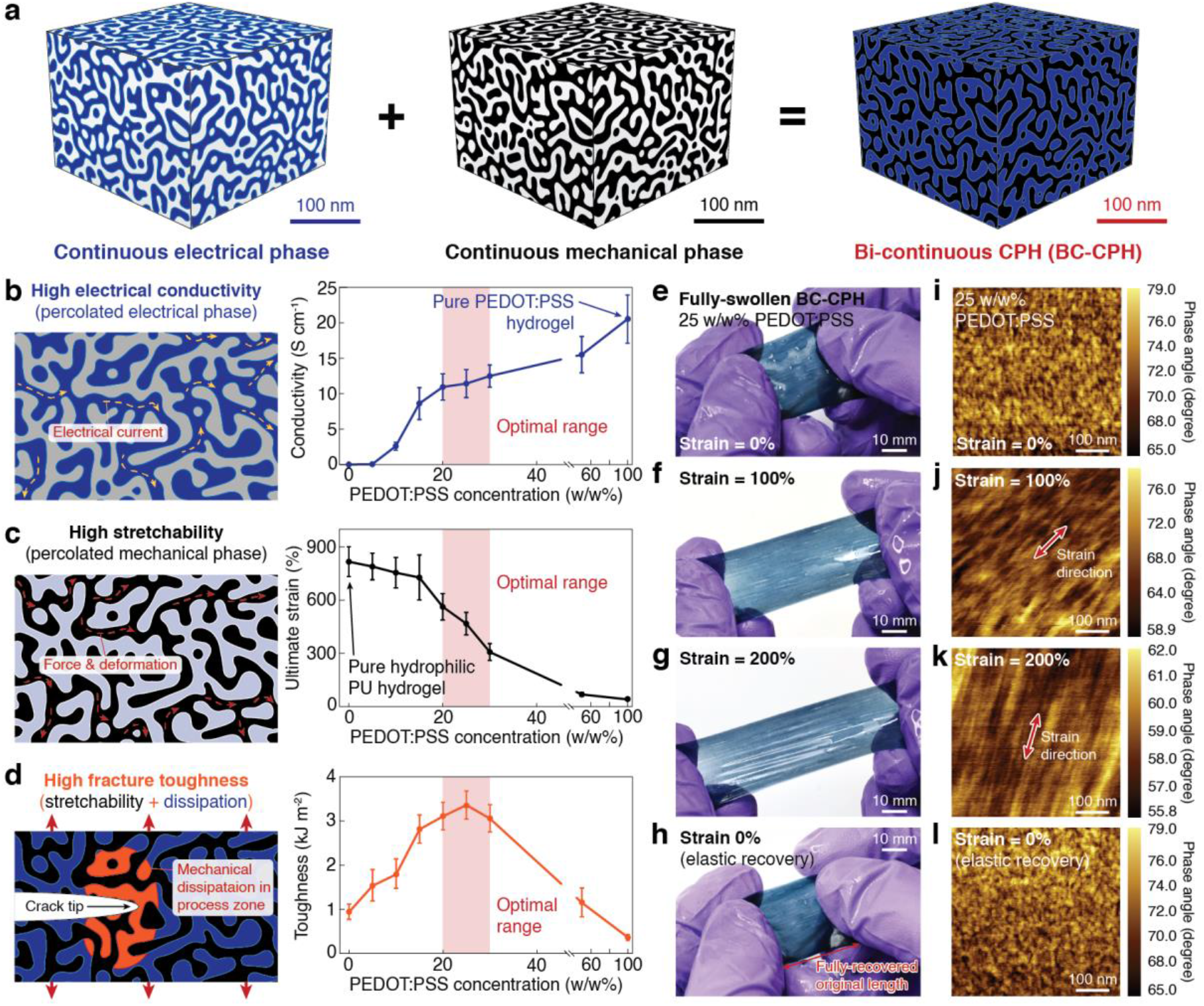
Design and implementation of the BC-CPH. **a**, Schematic illustrations of a bi-continuous conducting polymer hydrogel (BC-CPH) consisting of an electrical phase based on PEDOT:PSS and a mechanical phase based on hydrophilic polyurethane (PU). **b** to **d**, Schematic illustrations for mechanisms (left) and plots (right) for electrical conductivity (**b**), ultimate strain (**c**), and fracture toughness (**d**) vs. PEDOT:PSS concentration in the BC-CPH. **e** to **h**, Images of the fully-swollen BC-CPH with 25 w/w% PEDOT:PSS at engineering strain of 0% (**e**), 100% (**f**), 200% (**g**), and 0% after full elastic recovery (**h**). **i** to **l**, Corresponding AFM phase images of the BC-CPH with 25 w/w% PEDOT:PSS at engineering strain of 0% (**i**), 100% (**j**), 200% (**k**), and 0% after full elastic recovery (**l**). Error bars indicate SD; *N* = 4.

The mechanical phase is dominant in hydrogels with low PEDOT:PSS concentrations resulting in a low connectivity between electrical phases (**Fig. S4a**) and low electrical conductivity (**Fig. 1b**). Conversely, the electrical phase becomes dominant in hydrogels with high PEDOT:PSS concentrations resulting in a low connectivity between mechanical phases (**Fig. S4c**) and low stretchability (**Fig. 1c** and **Fig. S5**). We find that an optimal range of PEDOT:PSS concentration (20-30 w/w%) provides bi-continuous presence of both mechanical and electrical phases in the BC-CPH (**Fig. 1i** and **Fig. S4b**), simultaneously achieving high electrical conductivity (**Fig. 1b**) and stretchability (**Fig. 1c** and **Fig. S6**). The BC-CPH exhibits fully recoverable elastic deformation over 200% strain (**Fig. 1e** to **h**) maintaining its bi-continuous phases (**Fig. 1i** to **l**) without plastic deformation^20^.

Notably, the BC-CPH with the optimal range of PEDOT:PSS concentrations also exhibits a much higher fracture toughness over 3,000 J m^-2^ than mechanical or electrical phase-dominated hydrogels (**Fig. 1d** and **Fig. S7**). This marked enhancement in fracture toughness of the BC-CPH may originate from a similar mechanism to other tough hydrogels^9,10,21^, where the less stretchable PEDOT:PSS phase acts as a mechanical dissipater and the highly stretchable hydrophilic polyurethane phase maintains the integrity and elasticity of the BC-CPH (**Fig. 1d**).

The BC-CPH can provide favorable electrical and electrochemical properties including high electrical conductivity (over 11 S cm^-1^) even after over 5,000 cycles of 100% tensile strain (**Fig. 2a** and **b**), low impedance (**Fig. S8**), high charge storage capacity over 20 times higher than a Pt electrode (**Fig. 2d** and **e**), and high charge injection capacity over 3 times higher than a Pt electrode (**Fig. 2f** and **g**). The BC-CPH maintains favorable electrical and mechanical properties over 10,000 charging and discharging cycles (**Fig. 2e**), over 1M biphasic charge injections (**Fig. 2G**), and over 180 days of storage in physiological environments (**Fig. S9**). Notably, the BC-CPH exhibits strain-insensitive electrical resistance under moderate strain up to 50% (**Fig. 2c**), potentially due to dynamic aggregation of the electrical phase in the deformed state^22,23^ (**Fig. S10**).

**Fig. 2.**
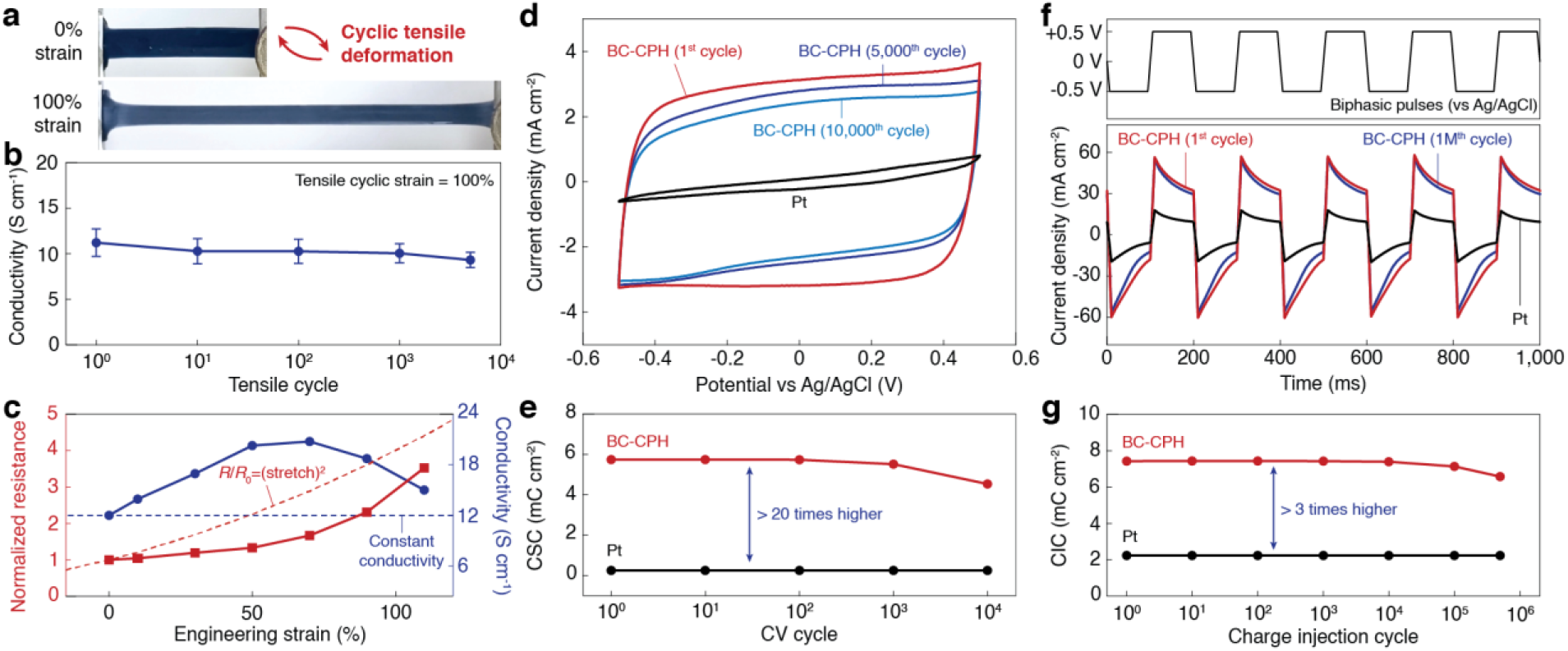
Electrical properties and stability of the BC-CPH. **a**, Images of the BC-CPH under cyclic tensile deformation of 100% engineering strain. **b**, Electrical conductivity vs. tensile cycle of the BC-CPH. **c**, Plots for electrical resistance normalized to the resistance of non-deformed state (*R*/*R*_0_, left axis) and electrical conductivity (right axis) vs. engineering strain of the BC-CPH. Stretch is engineering strain plus unity. **d**, Current density vs. potential plots for a Pt electrode and the BC-CPH at 1^st^, 5,000^th^, and 10,000^th^ cycle. **e**, Charge storage capacity (CSC) vs. cyclic voltammetry (CV) cycle for a Pt electrode and the BC-CPH. **f**, Biphasic input pulses (top) and the corresponding current density vs. time plots (bottom) for a Pt electrode and the BC-CPH at 1^st^ and 1M^th^ cycle. **g**, Charge injection capacity (CIC) vs. charge injection cycle for a Pt electrode and the BC-CPH. All tests were done in PBS. Error bars indicate SD; *N* = 4.

The design of BC-CPH allows its simple and facile fabrication from a viscous ink, which is readily applicable to various fabrication methods. The viscosity of the BC-CPH ink can be easily tuned by controlling the amount of solvent (70% ethanol) in the ink (**Fig. 3a**) without affecting the electrical or mechanical properties of the resultant BC-CPH (**Fig. 3b** and **c**). The low-viscosity BC-CPH ink can be used in various manufacturing methods including spin-coating^18^ (**Fig. 3d**) and electrospinning^24^ (**Fig. 3e**). The high-viscosity BC-CPH ink exhibits favorable rheological properties as a moldable and printable material, allowing fabrication of BC-CPH microstructures by micro-molding based on soft lithography (**Fig. 3f**) ^25^ as well as by 3D printing^15,26,27^ (**Fig. 3g**).

**Fig. 3.**
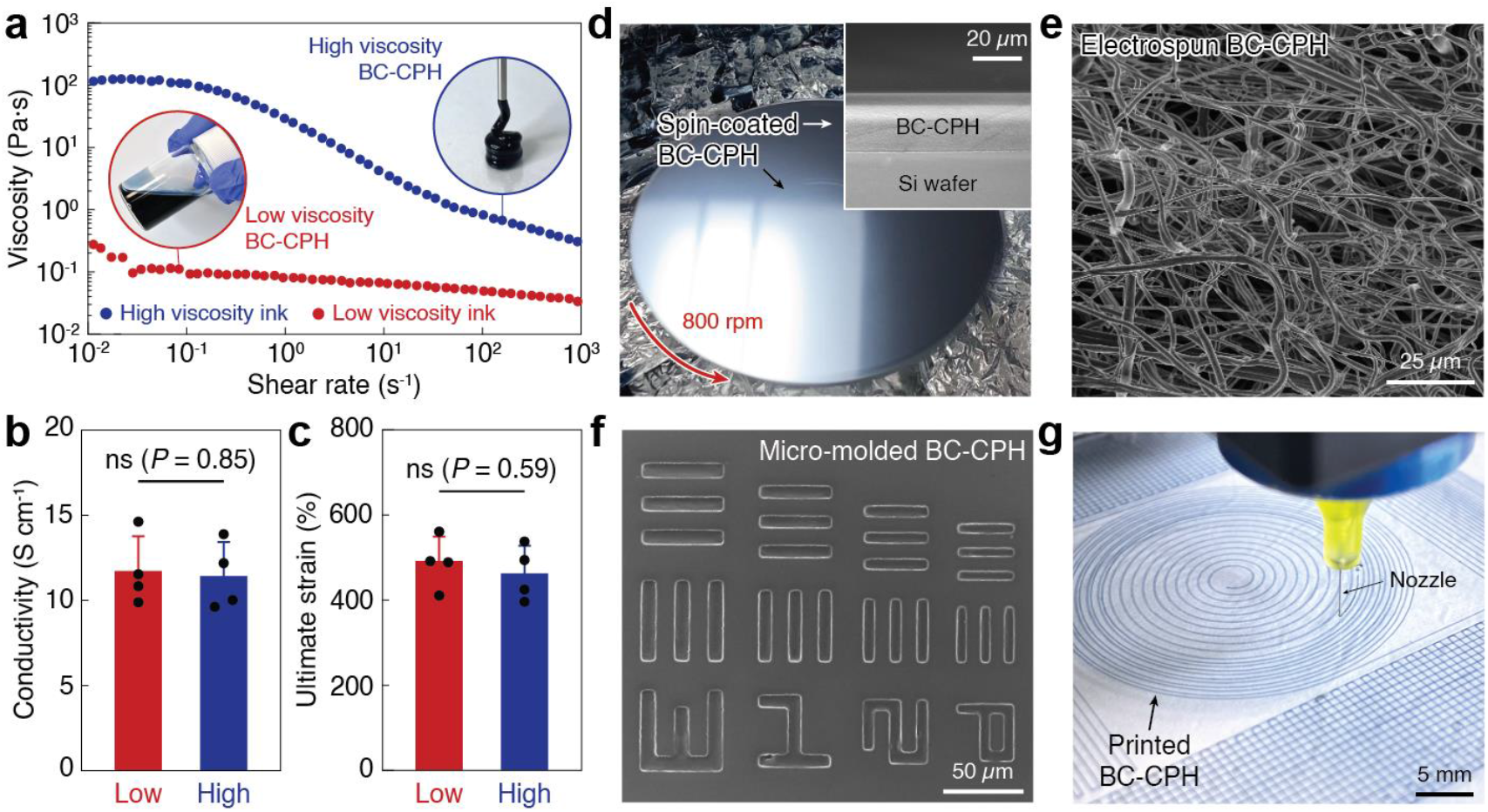
Applicability to diverse fabrication methods. **a**, Viscosity vs. shear rate plots for the BC-CPH inks with high and low viscosity. Inset figures show the high viscosity (inside blue circle) and low viscosity (inside red circle) BC-CPH inks. **b** and **c**, Electrical conductivity (**b**) and ultimate strain (**c**) of the BC-CPH prepared from high and low viscosity inks. **d**, Image of the spin-coated low viscosity BC-CPH ink. Top-right inset SEM image is the cross-sectional view of the spin-coated BC-CPH on a silicon wafer. **e**, SEM image of the electrospun low viscosity BC-CPH ink. **f**, SEM image of the micro-molded high viscosity BC-CPH ink from a SU-8 soft lithography mold. **g**, Image of the 3D-printed high viscosity BC-CPH ink. Error bars indicate SD; *N* = 4. Statistical significance is determined by two-sided Student t-test; ns, not significant.

Taking advantage of the BC-CPH’s ready-applicability to multi-material 3D printing, we demonstrate printing-based fabrication of hydrogel bioelectronic interfaces (**Fig. 4A**). In combination with printable bioadhesive (**Figs. S11a** and **S12**) and insulating (**Figs. S11b** and **S13**) hydrogel inks, the BC-CPH enables multi-material printing-based fabrication of hydrogel bioelectronic interfaces with tissue-like softness and water contents^28,29^ (**Fig. 4b**) in less than 10 min (**Fig. S14** and **movie S1**). The printed bioelectronic interfaces take the form of monolithic hydrogels (electrodes by the BC-CPH, encapsulation by the insulating hydrogel, and bio-integration by the bioadhesive hydrogel) with high flexibility in physiological environments (**Fig. 4c** and **Fig. S15**).

**Fig. 4.**
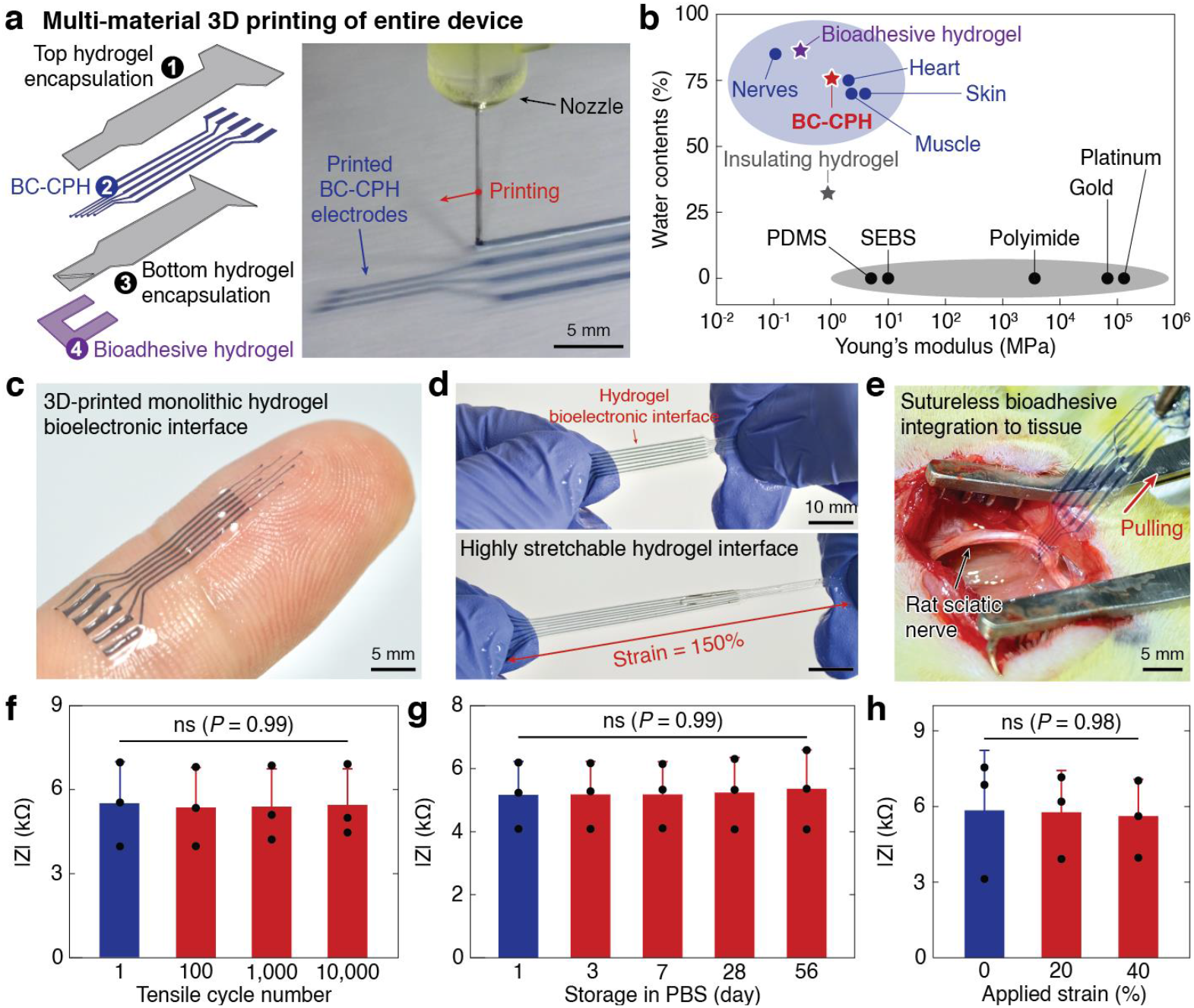
Hydrogel bioelectronic interface. **a**, Schematic illustration (left) and image (right) of multi-material 3D printing-based fabrication of the entire hydrogel bioelectronic interface. **b**, Young’s moduli vs. water contents for the hydrogel bioelectronic interface (BC-CPH, insulating and bioadhesive hydrogels), conventional electrode and encapsulation materials, and biological tissues ^29^. **c**, Image of a 3D-printed monolithic hydrogel bioelectronic interface on a fingertip. **d**, Images of the highly stretchable hydrogel bioelectronic interface before (top) and after (bottom) deformation. **e**, Image of sutureless robust bioadhesive integration of the hydrogel bioelectronic interface on a rat sciatic nerve. **f** to **h**, Impedance of one electrode channel in the hydrogel bioelectronic interface at 1 kHz under varying tensile cycle (**f**), storage time in a phosphate-buffered saline (PBS) bath at 37 °C (**g**), and tensile strain (**h**). Error bars indicate SD; *N* = 3. Statistical significance is determined by one-way ANOVA followed by Bonferroni’s multiple comparison test; ns, not significant.

Owing to the high stretchability of the BC-CPH and other constituent hydrogels (**Figs. S6** and **S16**), the hydrogel bioelectronic interfaces can withstand over 150% strain without failure (**Fig. 4d**). The bioadhesive in the hydrogel bioelectronic interfaces further provides rapid, robust, and sutureless integration to the target tissues^30–32^ (**Fig. 4e**, **Fig. S17**, and **movie S2**). The hydrogel bioelectronic interfaces maintain stable single-electrode impedance around 5 kΩ over 10,000 cycles of 20% tensile strains (**Fig. 4f**, **Figs. S18** and **S19**) and over 56 days of storage in physiological environments (**Fig. 4g** and **Fig. S19**). Taking advantage of the BC-CPH’s strain-insensitive electrical resistance at moderate deformation (**Fig. 2c**), the hydrogel bioelectronic interfaces also exhibit stable impedance up to 40%tensile strain (**Fig. 4h** and **Fig. S19c**), which can be highly favorable for bioelectronic interfaces in dynamic physiological environments^6,27^.

We conduct electrophysiological recording of rat hearts (**Fig. 5a** to **f**) and stimulation of rat sciatic nerves (**Fig. 5g** to **n**) and spinal cords (**Fig. S20**) by the hydrogel bioelectronic interfaces to demonstrate their long-term *in vivo* bioelectronic interfacing capability. Multi-material 3D printing allows a flexible choice of designs and fast manufacturing of the hydrogel bioelectronic interfaces for various target organs (**Fig. S21**). Favorable electrical properties of the BC-CPH electrodes in the hydrogel bioelectronic interfaces provide successful *in vivo* electrophysiological recording of rat hearts (epicardial signals, **Fig. 5d**) and stimulation of sciatic nerves (hindlimb movements, **Fig. 5j** and **l**) and rat spinal cords (forelimb movements, **Fig. S20d** and **f**) on day 0 post-implantation.

**Fig. 5.**
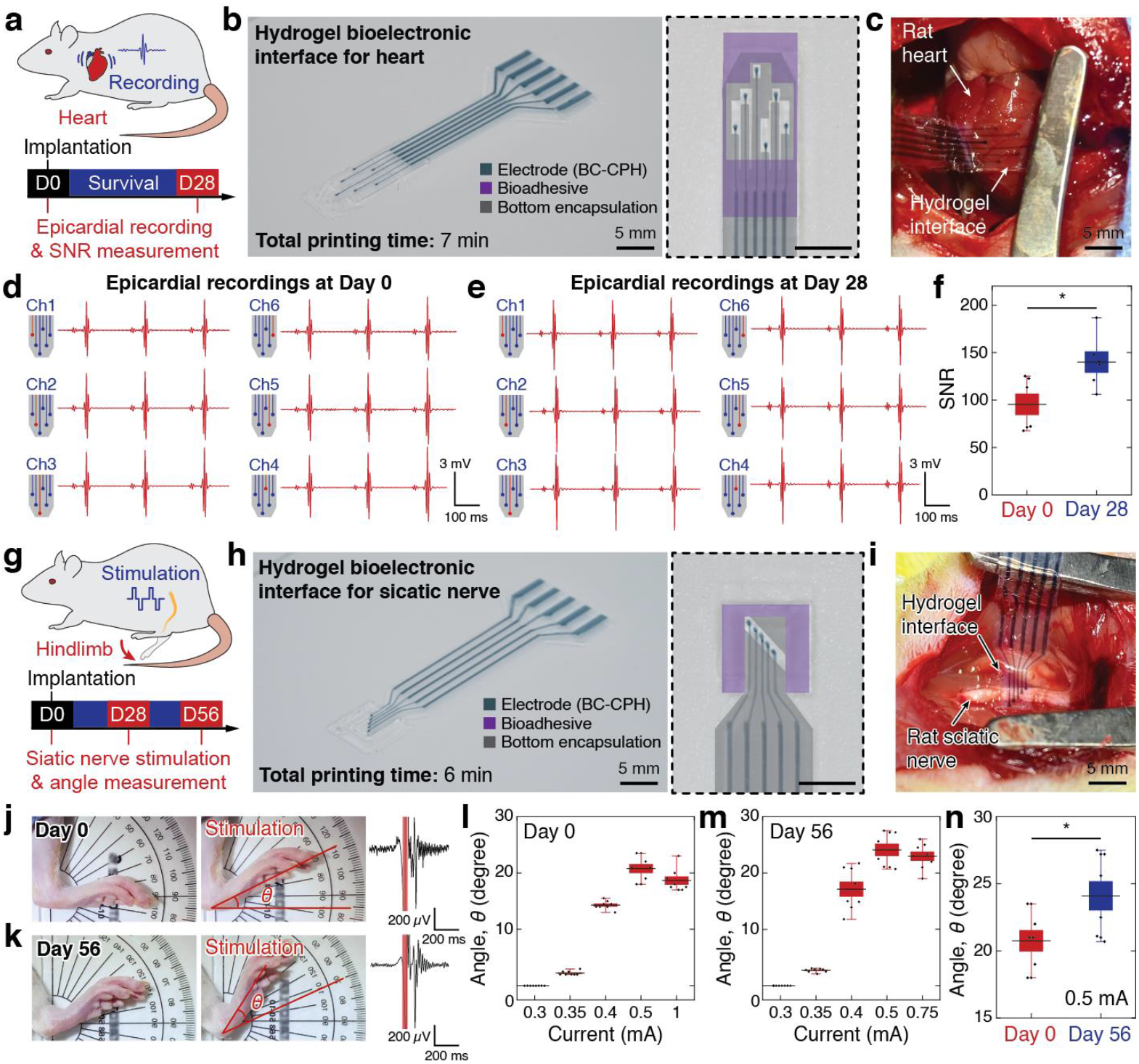
In vivo electrophysiological recording and stimulation. **a** and **g**, Schematic illustrations for rat heart recording (**a**) sciatic nerve stimulation (**g**) by the hydrogel bioelectronic interface. **b** and **h**, Images of the printed hydrogel bioelectronic interface for heart (**b**) and sciatic nerve (**h**) in the overall view (left) and the magnified view of electrodes (right). Different materials are marked with color overlays in the magnified view. **c** and **i**, Images of the implanted hydrogel bioelectronic interface on rat heart (**c**) and sciatic nerve (**i**). **d** and **e**, Epicardial recordings by different channels in the hydrogel bioelectronic interface on day 0 (**d**) and day 28 (**e**) post-implantation. **f**, Comparison of the signal-to-noise ratio (SNR) for epicardial recordings on day 0 and day 28 post-implantation. **j** and **k**, Images of rat hindlimb before (left) and after (middle) electrophysiological stimulation of the sciatic nerve by the hydrogel bioelectronic interface with corresponding EMG recordings (right) on day 0 (**j**) and day 56 (**k**) post-implantation. The red-shaded regions in the EMG recordings indicate the stimulation pulses. **l** and **m**, Rat hindlimb movement angles upon sciatic nerve stimulations by the hydrogel bioelectronic interface at varying stimulation currents on day 0 (**l**) and day 56 (**m**) post-implantation. **n**, Comparison of the rat hindlimb movement angles on day 0 and day 56 post-implantation with stimulation current of 0.5 mA. In box plots, center lines represent mean, box limits delineate standard error (SE), and whiskers reflect 5th and 95th percentile; *N* = 6-8. Statistical significance is determined by two-sided Student t-test; * *P* < 0.05.

Owing to their tissue-like properties and stability in physiological environments, the hydrogel bioelectronic interfaces exhibit stable integration to the target tissues (**Fig. S22**) and favorable tissue response during long-term *in vivo* implantation over 2 months (**Fig. 6**). Histological analysis by a blinded pathologist indicates that the hydrogel bioelectronic interfaces elicit very mild inflammation to the target tissues (**Fig. 6a** to **c**) with fibrotic tissues around the hydrogel bioelectronic interfaces significantly thinner than those around elastomer-based control devices (i.e., devices with polydimethylsiloxane (PDMS)-based encapsulation) and comparable to the sham group (**Fig. 6d**).

**Fig. 6.**
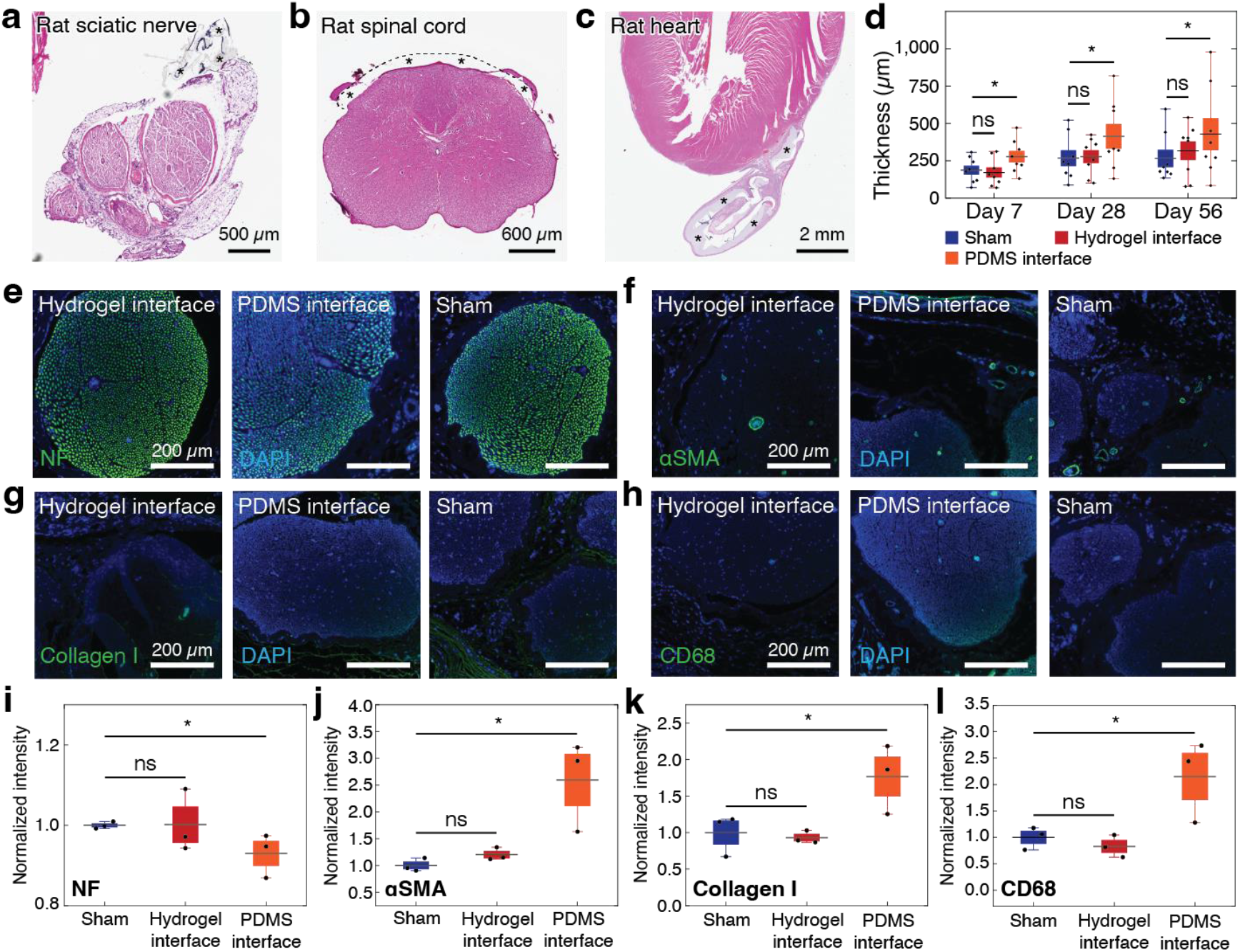
In vivo biocompatibility. **a** to **c**, Representative histological images of rat sciatic nerve (**a**), spinal cord (**b**), and heart (**c**) stained with hematoxylin and eosin (H&E) on day 28 post-implantation of the hydrogel bioelectronic interface. * indicates the implanted hydrogel bioelectronic interface. **d**, Sciatic nerve epineurium thickness measured from histological images on day 7, day 28, and day 56 post-implantation of the hydrogel bioelectronic interface, PDMS interface, and sham group (no device implantation). **e** to **h**, Representative immunofluorescence images of rat sciatic nerve on day 28 post-implantation of the hydrogel bioelectronic interface, PDMS interface, and sham group (no device implantation). Cell nuclei are stained with DAPI (blue). Green fluorescence corresponds to the expression of neurofilament (NF, **e**), fibroblasts (αSMA, **f**), collagen (collagen I, **g**), and macrophages (CD68, **h**), respectively. **i** to **l**, Normalized fluorescence intensity plots for the expression of NF (**i**), αSMA (**j**), Collagen I (**k**), and CD68 (**l**) in different groups. In box plots, center lines represent mean, box limits delineate standard error (SE), and whiskers reflect 5th and 95th percentile; *N* = 8 (d); *N* = 3 (i to l). Statistical significance is determined by two-sided Student t-test; ns, not significant; * *P* < 0.05.

We further conduct immunofluorescence analysis for various markers to evaluate tissue damage such as neurofilament (NF) and foreign body response including fibroblasts (αSMA), macrophages (CD68), collagen (Collagen I), and T cells (CD3) (**Figs. 6e** to **l, S23** to **S26**). The quantitative analysis of fluorescence intensity indicates that the hydrogel bioelectronic interfaces elicit comparable expression of all markers to the sham group on day 7, 28, and 56 post-implantations on rat sciatic nerves (**Figs. 6i** to **l, S23** to **S24**). In contrast, the elastomer-based control devices induce significantly lower expression of NF and higher expression of αSMA, Collagen I, and CD68 compared to the sham group on day 28 post-implantation, indicating potential damage of neural tissues and higher foreign body response (**Fig. 6i** to **l**).

Enabled by the favorable *in vivo* biocompatibility and stability, the hydrogel bioelectronic interfaces can provide stable long-term electrophysiological recording and stimulation of rat hearts (**Fig. 5e**), sciatic nerves (**Fig. 5k** and **m**), and spinal cords (**Fig. S20e** and **g**). Notably, the efficacy of electrophysiological recording and stimulation by the hydrogel bioelectronic interfaces increases in the longer term compared to the efficacy on day 0 (i.e., right after implantation), demonstrated by significantly higher signal-to-noise ratio (SNR) of the measured epicardial signals for heart recording for all six electrode channels (**Fig. 5f**) and by significantly higher hindlimb joint angle movement for sciatic nerve stimulation (**Figs. 5n** and **S27**) and forelimb movement distance for spinal cord stimulation (**Fig. S20h**) for the same injected current. This enhancement in long-term electrophysiological efficacy might be facilitated by the tissue-like properties, atraumatic bioadhesive integration, and resultant favorable tissue interaction of the hydrogel bioelectronic interfaces^31,33^.

By addressing the lingering challenges in conductive hydrogels, the BC-CPH provides a promising material for tissue-like bioelectronics. Enabled by the unique set of advantages of the BC-CPH, we 3D print monolithic all-hydrogel bioelectronic interfaces capable of long-term high-efficacy electrophysiological stimulation and recording of diverse tissues and organs in rat models. This work may offer a versatile tool and platform toward a vision of hydrogel bioelectronics^1^ and better electrical interfacing between machines and biological systems.

## Acknowledgments

The authors thank the Koch Institute Swanson Biotechnology Centre, K. Cormier, and the Histology Core for the technical support and histological processing, Dr. R. Bronson at Harvard Medical School for the histological evaluations.

## Funding

This work is supported by National Institute of Health (1-R01-HL153857-01) and Massachusetts Institute of Technology. B.L. acknowledges the support from Jiangxi Science and Technology Normal University. H.Y. acknowledges the support from Samsung Scholarship.

## Author contributions

H.Y., B.L. and X.Z. developed the concept and materials for the BC-CPH. H.Y. and T.Z. developed the materials and method for the printing-based fabrication and application of the hydrogel bioelectronic interface. B.L., H.Y., F.H., F.T., and J.X. conducted the electrical and mechanical characterizations of the BC-CPH. T.Z. and H.Y. conducted the electrical and mechanical characterizations of the hydrogel bioelectronic interface. T.Z., H.Y., and J.W. designed and conducted the *in vivo* animal studies. H.Y. and H.R. developed the materials and method for the printable bioadhesive. Z.S. and G.G. conducted the AFM phase imaging. H.Y. and X.Z. developed the multi-material printing platform. X.Z. initiated and supervised the research on hydrogel bioelectronics. H.Y., T.Z., and X.Z. wrote the manuscript with inputs from all co-authors.

## Competing interests

T.Z., H.Y., B.L. and X.Z are inventors of a patent application on the BC-CPH filed by Massachusetts Institute of Technology. Other authors declare no competing interests.

## Data and materials availability

All data is available in the main text or the supplementary materials.

## Supplementary Materials

Materials and Methods

Supplementary Text

Figs. S1 to S27

Movies S1 to S2

References (*34-44*)

## Notes

### Competing Interest Statement

The authors have declared no competing interest.

