## Supplementary Materials for "3D Printable High Performance Conducting Polymer Hydrogel for All-Hydrogel Bioelectronics"

**This PDF file includes:**

Materials and Methods
Supplementary Text
Figs. S1 to S27
Captions for Movies S1 to S2
References

**Other Supplementary Materials for this manuscript include the following:**

Movies S1 to S2

### Materials and Methods

Materials

For the preparation of the BC-CPH, aqueous poly(3,4-ethylenedioxythiophene):polystyrene sulfonate (PEDOT:PSS) suspension (Clevios$^{TM}$ PH1000; Heraeus Electronic Materials), dimethyl sulfoxide (DMSO; Sigma-Aldrich), ethanol (Sigma-Aldrich), and hydrophilic polyurethane (HydroMed D3; AdvanSource Biomaterials) were used. For the preparation of the bioadhesive hydrogel ink, acrylic acid (Sigma-Aldrich), hydrophilic polyurethane (HydroMed D3, AdvanSource Biomaterial), benzophenone (Sigma-Aldrich), α-ketoglutaric acid (Sigma-Aldrich), 1-Ethyl-3-(3-dimethylaminopropyl) carbodiimide (EDC; Sigma-Aldrich), and *N*-Hydroxysuccinimide (NHS; Sigma-Aldrich) were used. For the preparation of the insulating hydrogel ink, low water contents hydrophilic PU (HydroThane AL-25 80A, AdvanSource Biomaterials), dimethylformamide (DMF; Sigma-Aldrich), and tetrahydrofuran (THF; Sigma-Aldrich) were used. For printing of the hydrogel bioelectronic interface, poly (vinyl alcohol) (PVA; Mw 31,000-50,000, 87%-89% hydrolyzed, Sigma-Aldrich), 100-µm and 200-µm nozzles (Nordson EFD), and 5-mL syringe barrel (Nordson EFD) were used.

Methods

**Preparation of the BC-CPH.** An aqueous PEDOT:PSS suspension was stirred vigorously for 6 h at room temperature and filtered with a syringe filter (0.45-µm polypropylene). The filtered PEDOT:PSS suspension was transferred to a clean glass vial and cryogenically frozen by submerging in a liquid nitrogen bath. The cryogenically frozen PEDOT:PSS suspension was lyophilized for 72 h to isolate PEDOT:PSS nanofibrils. The isolated PEDOT:PSS nanofibrils were re-dispersed with deionized water and DMSO mixture (water:DMSO = 85:15 v/v) at the concentration of 7 w/w%, followed by thorough mixing and homogenization by a mortar grinder (RM 200; Retch). The re-dispersed PEDOT:PSS suspension was then mixed with 7 w/w% hydrophilic polyurethane in ethanol solution (ethanol:deionized water = 95:5 v/v) at varying ratio. The mixture was further diluted with 70% ethanol solution to prepare a high viscosity printable BC-CPH ink (7 w/w% polymer concentration) or a low viscosity spin-coatable BC-CPH ink (0.5 w/w% polymer concentration), followed by filtering with a syringe filter (10-µm polypropylene). The BC-CPH was prepared by air drying the BC-CPH ink at room temperature for 24 h and swelling the dried sample in a large volume of phosphate-buffered saline (PBS). Otherwise mentioned, the BC-CPH with 25 w/w% PEDOT:PSS concentration (PEDOT:PSS : hydrophilic polyurethane = 1:3 w/w) was used.

**Preparation of the bioadhesive hydrogel ink.** To prepare a precursor solution, add 32 w/w% acrylic acid, 8 w/w% hydrophilic PU, 1.1 w/w% benzophenone, and 0.1 w/w% α-ketoglutaric acid in ethanol solution (ethanol:deionized water = 2:1 v/v). To graft polyacrylic acid to the hydrophilic PU (PU-PAA), the homogeneously mixed precursor solution was transferred to a sealed glass vial and cured in an ultraviolet (UV) crosslinker (365 nm, 15 W powder) for 120 min. The cured precursor solution was then purified by using cellulose dialysis bags in a pure ethanol bath for 24 h (bath replaced every 12 h) followed by in a deionized water bath for 24 h (bath replaced every 12 h) with continuous magnetic stirring. The purified PU-PAA samples were cut into small pieces and thoroughly dried in a desiccating oven at 70 °C for 24 h. To prepare the bioadhesive hydrogel ink, the dried PU-PAA was dissolved in 70 % ethanol solution at the concentration of 20 w/w%. 2 w/w% EDC and 2 w/w% NHS were added to the bioadhesive hydrogel ink before printing to introduce NHS ester groups to PU-PAA. The bioadhesive was prepared by air drying the printed ink at room temperature for 24 h and used in the dry state to facilitate wet adhesion. To prepare the bioadhesive hydrogel in mechanical characterizations, the dry bioadhesive sample was swollen in a large volume of PBS.

**Preparation of the insulating hydrogel ink.** To prepare the insulating hydrogel ink, low water contents hydrophilic PU was dissolved in a solvent mixture (DMF:THF = 1:1 v/v) at the concentration of 25 w/w%. The insulating hydrogel was prepared by air drying the printed ink at 70 °C for 3 h and swelling the dried sample in a large volume of PBS.

**Mechanical characterizations.** All mechanical characterizations were performed by using the fully swollen samples in PBS. Mechanical properties of the samples were characterized by a mechanical testing machine (U-Stretch with 4.4 N load cell; CellScale). All mechanical characterizations were performed within the submersion stage filled with PBS to avoid dehydration of the sample at a constant crosshead speed of 50 mm min$^{-1}$.

For measurement of ultimate strain and Young's modulus, dog-bone samples (10 mm in gauge length; 3 mm in width; 0.2 mm in thickness) were used. The ultimate strain of the sample was measured based on the engineering strain at which the sample ruptured. Young's modulus of the sample was measured by fitting the engineering stress vs. engineering strain curve with the incompressible neo-Hookean model for uniaxial extension,

$$S = \frac{E}{3}\left(\varepsilon + 1 - \frac{1}{(\varepsilon + 1)^2}\right)$$

where $S$ is the engineering stress, $E$ is the Young's modulus of the sample, and $\varepsilon$ is the engineering strain.

For measurement of fracture toughness, rectangular samples (20 mm in length; 40 mm in width; 0.2 mm in thickness) without or with notch (10 mm in length) were used. The fracture toughness of the sample was calculated by following the previously reported method based on tensile tests of unnotched and notched samples [10].

For measurement of interfacial toughness, dry bioadhesives (30 mm length, 10 mm in width, 0.2 mm in thickness) adhered to various tissues (sciatic nerve, spinal cord, heart, muscle) were tested by the 180-degree peel test [30]. The measured force reached a plateau as the peeling process entered the steady-state. Interfacial toughness was determined by dividing two times the plateau force by the width of the sample. Poly (methyl methacrylate) films (with a thickness of 50 μm; Goodfellow) was applied using cyanoacrylate glue (Krazy Glue) as a stiff backing for the tissues and hydrogels. All rat tissues used for the measurement of interfacial toughness were collected and used following the protocol reviewed and approved by the Committee on Animal Care at the Massachusetts Institute of Technology.

Rheological characterizations of the various inks were conducted by using a rotational rheometer (AR-G2; TA Instrument) with 20-mm diameter steel parallel-plate geometry. Viscosity was measured as a function shear rate by steady-state flow tests with a logarithmic sweep of shear rate (0.01 to 1,000 $s^{-1}$). All rheological characterizations were conducted at 25 °C with a preliminary equilibration time of 1 min.

**Electrical characterizations.** For electrical characterizations, the free-standing BC-CPH films (30 mm in length; 5 mm in width; 0.1 mm in thickness) or the hydrogel bioelectronic interfaces for sciatic nerve fully swollen in PBS were used. The electrical conductivity of the sample was measured by using a standard four-point probe (Keithley 2700; Keithley). Pt wire electrodes (0.5 mm in diameter) were attached to the surface of the sample by applying the silver paste. The electrical conductivity of the samples was calculated as

$$\sigma = \frac{I \times L}{V \times W \times T}$$

where σ is the electrical conductivity, $I$ is the current flowing through the sample, $L$ is the distance between the two electrodes for voltage measurement, $V$ is the voltage across the electrodes, $W$ is the width of the sample, and $T$ is the thickness of the sample.

Electrochemical impedance spectroscopy (EIS) measurements of the sample were performed by using a potentiostat/galvanostat (1287A, Solartron Analytical) and a frequency response analyzer (1260A, Solatron Analytical) in an electrochemical cell installed with the sample as a working electrode, a Pt sheet as a counter electrode, an Ag/AgCl wire as a reference electrode, and PBS as an electrolyte. The frequency range between 0.1 and 100 kHz was scanned with an applied bias of 0.01 V vs. Ag/AgCl.

Cyclic voltammetry (CV) measurements were performed by using a potentiostat/galvanostat (VersaSTAT 3; Princeton Applied Research) with the potential window of ± 0.5 V vs. Ag/AgCl and the potential scan rate of 150 mV $s^{-1}$ in an electrochemical cell installed with the sample as a working electrode, a Pt sheet as a counter electrode, an Ag/AgCl wire as a reference electrode, and PBS as an electrolyte. The CSC of the sample was calculated from the measured CV data as

$$\mathrm{CSC} = \int_{E_2}^{E_1} \frac{i(E)}{2vA} dE$$

where $v$ is the scan rate, $E_2$ and $E_1$ are the potential window, $i$ is the current at each potential, and $A$ is the area of the sample.

To measure charge injection capacity (CIC), biphasic pulses at 200 ms ± 0.5 V were applied by using a multi-channel potentiostat (VMP3, Bio-Logic Science Instruments) in an electrochemical cell installed with the sample as a working electrode, a Pt sheet as a counter electrode, an Ag/AgCl wire as a reference electrode, and PBS as an electrolyte. The CIC of the sample was calculated from the measured output voltage and current as

$$\mathrm{CIC} = \frac{Q_{\mathrm{inj(c)}} + Q_{\mathrm{inj(a)}}}{A}$$

Where $Q_{inj(c)}$ is the total delivered (or injected) charge in the cathodal phase, $Q_{inj(a)}$ is the total delivered (or injected) charge in the anodal phase, and $A$ is the area of the sample.

**AFM phase imaging.** Atomic force microscope (AFM) phase images were acquired by atomic force microscope (MFP-3D, Asylum Research). Undeformed or stretched free-standing BC-CPH films were directly attached to the sample stage by double-sided carbon tape.

**SEM imaging.** Scanning electron microscope (SEM) images of samples were taken by a SEM facility (Zeiss Supra 40; Zeiss) with 5 nm gold sputtering to enhance image contrasts.

**Micro-molding by soft lithography.** A 3-inch silicon wafer (University Wafer, Inc.) was cleaned by oxygen plasma (50 W) for 1 min. Photoresist SU-8 (SU-8 2010; MicroChem) was spin-coated on the wafer at 2,000 rpm for 1 min, followed by pre-baking sequentially at 60 °C for 1 min and 95 °C for 4 min. The photoresist was then patterned by photolithography with a mask aligner (SUSS MA6 mask aligner; SUSS MicroTec). After the photolithography exposure, the silicon wafer was post-baked sequentially at 65 °C for 1 min and 95 °C for 4 min. The SU-8 photoresist was developed (SU-8 Developer; MicroChem) for 1.5 min, followed by rinsing with isopropanol and drying with nitrogen gas. The high viscosity BC-CPH ink was applied on the prepared mold, dried for 30 min at 40 °C and peeled off from the substrate.

**Printing of the hydrogel bioelectronic interface.** Before the printing of the hydrogel bioelectronic interface, a layer of PVA was introduced as a water-dissolvable substrate for the hydrogel interface. To introduce the PVA layer, an aqueous PVA solution (30 w/w% in deionized water) was spin-coated on a printing substrate at 600 rpm for 1 min followed by drying at 70 °C for 1 h. Multi-material printing was conducted by a custom-designed 3D printer based on a Cartesian gantry system (AGS1000; Aerotech) with various sizes of nozzles (200- and 100-µm nozzles) connected to syringe barrels loaded with the BC-CPH, bioadhesive, and insulating hydrogel inks [26]. Printing paths were prepared by CAD drawings (SolidWorks; Dassault Systèmes) and converted into G-codes by a commercial software package (CADFusion; Aerotech) to command the X-Y-Z motion of the printer head. All hydrogel bioelectronic interfaces for animal studies were prepared in an aseptic manner and were further disinfected under UV light for 15 min.

**Vertebrate animal subjects.** Female Sprague Dawley rats (225–250 g, Strain Code 400, Charles River) were used in this work. All animal studies were reviewed and approved by the Committee on Animal Care at the Massachusetts Institute of Technology. The animal care and use programs at Massachusetts Institute of Technology meet the requirements of the Federal Law (89-544 and 91-579) and NIH regulations and are also accredited by the American Association for Accreditation of Laboratory Animal Care (AAALAC).

***In vivo* sciatic nerve surgeries.** Animals were anesthetized using 3% inhaled Isoflurane. Animals were placed in the prone position over a heating pad for the duration of the surgery. Anesthesia was maintained with a nose cone and 1-2% Isoflurane in $O_2$. Respiratory rate and quality were used to monitor the depth of anesthesia. Sterile eye ointment was applied after anesthesia and before shaving to minimize the risk of corneal irritation, dehydration, and sensitization during surgical procedures. Before starting the surgery, the depth of anesthesia was checked by monitoring of tail/toe pinch response. The surgical sites of the animals were shaved to remove dorsocaudal region hair, and the shaved area was prepared with an application of Betadine and three subsequent applications of 70% ethanol rinses, each with a contact time of at least two minutes. A 2-cm incision was made through the dermis of the animal's hindlimb, exposing the subcutaneous tissue. The sciatic nerve was exposed by separating muscles close to the femur. The hydrogel bioelectronic interface or the PDMS interface (control) were implanted on the surface of the exposed sciatic nerve. For the sham group, no device was implanted. The incision was closed with 4–0 sutures and 3–6 ml of warm saline was injected subcutaneously as post-surgical hydration support.

***In vivo* spinal cord surgeries.** Animals were anesthetized using 3% inhaled Isoflurane. Animals were placed in the prone position over a heating pad for the duration of the surgery. Anesthesia was maintained with a nose cone and 1-2% Isoflurane in $O_2$. Respiratory rate and quality were used to monitor the depth of anesthesia. Sterile eye ointment was applied after anesthesia and before shaving to minimize the risk of corneal irritation, dehydration, and sensitization during surgical procedures. Before starting the surgery, the depth of anesthesia was checked by monitoring of tail/toe pinch response. The surgical site of the animals was shaved to remove back the hair from slightly rostral to ears to the middle of the animal's back. The shaved area was prepared with an application of Betadine and three subsequent applications of 70% ethanol rinses, each with a contact time of at least two minutes. A small incision around 10 mm in length above the vertebrae of interest (C4-C6) was created by using a scalpel blade. The size of the opening in the skin was then increased by blunt dissection with forceps or surgical scissors. Further incisions were made through the muscle layers over the spinal column using a scalpel blade. The surgical field was made by retracing the muscle tissues by using a sterile autoclaved soft tissue retractor. All overlying tissues from the dorsal laminae were removed by using a spring scissor and sterile cotton swabs, and the spinal column was secured with rat-toothed forceps. A laminectomy was conducted by grabbing the entire lamina with a rongeur and slowly breaking the lamina with a rongeur or spring scissors. The broken pieces of the spine were gently pulled upwards and the surrounding connective tissues were cleaned off to expose the spinal cord. The hydrogel bioelectronic interface was implanted on the spinal cord epidurally from the entry point with help of a sterilized thin polyethylene terephthalate film (100 µm-thick, Goodfellow). For the sham group, no device was implanted. The incision was closed with 4–0 sutures and 3–6 ml of warm saline was injected subcutaneously as post-surgical hydration support.

***In vivo* heart surgeries.** Animals were anesthetized using 3% inhaled Isoflurane. Anesthesia was maintained with a nose cone and 1-2% Isoflurane in $O_2$. Respiratory rate and quality were used to monitor the depth of anesthesia. Sterile eye ointment was applied after anesthesia and before shaving to minimize the risk of corneal irritation, dehydration, and sensitization during surgical procedures. Before starting the surgery, the depth of anesthesia was checked by monitoring of tail/toe pinch response. Chest hair was then removed. Endotracheal intubation was performed, and the animals were connected to a mechanical ventilator (Model 683, Harvard Apparatus) and placed supine over a heating pad for the duration of the surgery. The shaved area was prepared with an application of Betadine and three subsequent applications of 70% ethanol rinses, each with a contact time of at least two minutes. The heart was exposed via a thoracotomy in the third or fourth left intercostal space and the pericardium was removed with fine forceps. The hydrogel bioelectronic interface was implanted on the epicardium of the exposed heart. For the sham group, no device was implanted. The incision was closed with 4–0 sutures and 3–6 ml of warm saline was injected subcutaneously as post-surgical hydration support. The animals were ventilated with 100% oxygen until autonomous breathing was regained, and the intubation catheter was removed.

***In vivo* sciatic nerve stimulation.** On day 0, day 28, and day 56 post-implantation, the implanted animals were anesthetized by using inhaled Isoflurane. Input/output (I/O) end of the implanted hydrogel interface was connected to a RHS Stim/Recording Controller (Intan Technologies) through a custom-designed PCB board with a flat flexible cable (Digi-Key Electronics). A needle electrode was inserted into the skin of the other hindlimb as the reference and ground. Cathode-first charge-balanced electrical pulses (1 Hz, 0.2–1 mA) with a width of 0.2 ms were applied by the RHS Stim/Recording Controller. A protractor marker was placed under the animal hindlimb to measure the change in the angle of the ankle joint. A Pt electrode (A-M Systems) was inserted into desired muscles for EMG recordings through the RHS Stim/Recording Controller and RHS amplifier (Intan Technologies) at a sampling rate of 20 kHz.

***In vivo* spinal cord stimulation.** On day 0 and day 28 post-implantation, the implanted animals were anesthetized by using inhaled Isoflurane. The animals were placed on a custom-made body supporter to allow forelimbs to move freely. Input/output (I/O) end of the implanted hydrogel interface was connected to the RHS Stim/Recording Controller through the custom-designed PCB board with the flat flexible cable. A needle electrode was inserted into the skin on the back as the reference and ground. Cathode-first charge-balanced electrical pulses (1 Hz, 0.3–2.5 mA) with a width of 0.2 ms were applied by the RHS Stim/Recording Controller. A ruler was placed between the animal forelimb and the custom-made body-supporter to measure the movement distance of animal forelimbs. A Pt electrode was inserted into desired muscles for EMG recordings through the RHS Stim/Recording Controller and RHS amplifier at a sampling rate of 20 kHz.

***In vivo* epicardial recording.** On day 0 and day 28 post-implantation, the implanted animals were anesthetized by using inhaled Isoflurane. Input/output (I/O) end of the implanted hydrogel interface was connected to the RHS Stim/Recording Controller through the custom-designed PCB board with the flat flexible cable. A needle electrode was inserted into the left forelimb as the reference and ground. Epicardial signals were then collected with the RHS Stim/Recording Controller and RHS amplifier at a sampling rate of 20 kHz. The signal-to-noise ratio (SNR) was calculated by dividing the average peak-to-peak amplitude of recorded signals by noise derived from noise estimation of corresponding recording traces.

**Histology**. At the end of each study, the animals were euthanized by $CO_2$ inhalation. The tissue of interest was excised and fixed in 10% formalin solution for 24 h for histological processing. The fixed tissue samples were placed in 70% ethanol and submitted for paraffin embedding, sectioning, and hematoxylin and eosin (H&E) staining at the Hope Babette Tang (1983) Histology Facility in the Koch Institute for Integrative Cancer Research at Massachusetts Institute of Technology. The completed histology slides were scanned by using a digital slide scanner (Aperio, Leica).

**Immunohistochemistry and immunofluorescence analysis.** For immunofluorescence analysis, the sectioned slides were deparaffinized and rehydrated in deionized water. Antigen retrieval was performed using a steam method during which the slides were steamed in IHC-Tek Epitope Retrieval Solution (IW-1100) for 35 min and then cooled for 20 min. Then the slides were washed in three changes of PBS for 5 min per cycle. After washing, the slides were incubated in primary antibodies (1:1000 Rabbit anti-neurofilament for neurofilament (ab8135, Abcam); 1:200 mouse anti-α-SMA for fibroblast (ab7817, Abcam); 1:200 mouse anti-CD68 for macrophages (ab201340, Abcam); 1:200 rabbit anti-collagen-I for collagen (ab21286, Abcam); 1:100 rabbit anti-CD3 for T cells (ab5690, Abcam)) diluted with IHC-Tek Antibody Diluent for 1 h at room temperature. The slides were then washed three times in PBS and incubated with Alexa Fluor 488 labeled anti-rabbit or anti-mouse secondary antibody (1:400, Jackson Immunoresearch) for 30 min. The slides were washed in PBS and then counterstained with DAPI for 20 min. A laser confocal microscope (SP 8, Leica) was used for image acquisition. MATLAB (version R2018b) was used to quantify the fluorescence intensity of expressed antibodies. All the images were transformed into double-precision images for analysis. Fluorescence intensities were calculated and normalized against the mean values of the corresponding sham groups. All analyses were blinded with respect to the experimental conditions.

**Statistical analysis.** Prism 9 (GraphPad, version 9.1) software was used to assess the statistical significance of all comparison studies in this work. Data distribution was assumed to be normal for all parametric tests, but not formally tested. In the statistical analysis for comparison between multiple samples, one-way ANOVA followed by Bonferroni's multiple comparison test was conducted with the threshold of * $P < 0.05$, ** $P \leq 0.01$, and *** $P \leq 0.001$. In the statistical analysis between two data groups, two-sided Student's t-test was used with the threshold of * $P < 0.05$, ** $P \leq 0.01$, and *** $P \leq 0.001$.

### Supplementary Text

Printable bioadhesive hydrogel for rapid sutureless wet adhesion to the target tissue

The recent advances in bioadhesives have enabled rapid and sutureless adhesion to wet biological tissues and organs. For example, the dry double-sided tape (DST) and its dry-crosslinking mechanism allow the formation of robust wet adhesion to various tissues in few seconds without the need of external stimuli such as UV light[30]. Such rapid, atraumatic, and preparation-free integration to wet tissues is highly advantageous to form a stable and conformal contact between the implanted bioelectronic interface and the target tissue[30,31]. However, the existing bioadhesives are typically prepared in the form of prefabricated films that require additional assembly procedures to combine with the bioelectronic device before or during implantation *in vivo*. In this work, we developed a printable bioadhesive ink that can be readily incorporated into multi-material printing of the hydrogel bioelectronic interface while providing the rapid, robust, and preparation-free adhesion capability to wet tissues comparable to that of the DST.

To synergistically achieve printability, bioadhesive capability, mechanical robustness in the swollen state, and compatibility to other components of the hydrogel bioelectronic interface (i.e., the BC-CPH, the insulating hydrogel), we covalently graft a bioadhesive polymer used in the DST (PAA-NHS ester) to hydrophilic polyurethane backbones by adopting the benzophenone-based radical grafting method[34-36]. The resultant bioadhesive can be dissolved in ethanol solution at a high concentration to yield a highly viscous printable ink (Fig. S11a) similar to the printable BC-CPH ink. The printed bioadhesive ink can be dried to provide a dry bioadhesive like the DST while forming robust mechanical integration with other components of the hydrogel bioelectronic interface (i.e., the BC-CPH, the insulating hydrogel) owing to the shared hydrophilic polyurethane-based composition of these materials.

The dry printed bioadhesive can provide rapid adhesion to wet tissues based on the dry-crosslinking mechanism[30,37]. Upon contacting the wet tissue surface, the negatively charged carboxylic acid groups in the PAA-NHS ester provide the quick hydration and swelling of the bioadhesive absorbing the interfacial water on the wet tissue surface. Simultaneously, the carboxylic acid groups in the bioadhesive physically crosslink with the tissue surface via hydrogen bonds, providing rapid initial adhesion (Fig. S12a). Subsequently, the NHS ester groups in the bioadhesive form covalent crosslinks with primary amine groups on the tissue surface via amide bonds within few minutes from the initial adhesion, providing a stable adhesion with the target tissue (Fig. S12b).

Printable insulating hydrogel for electrical encapsulation of the hydrogel interfaces

Hydrogels in the swollen state are ionically conductive due to dissolved ionic species within highly-hydrated regions of the hydrogel networks[3]. Since physiological environments are wet with abundant ionic species (e.g., dissolved salts and charged biomolecules), hydrogels with high water contents typically exhibit ionic conductivity similar to that of biological tissues in physiological environments [1,3]. Hydrogels' ionic conductivity is dependent on their water contents as the ionic conductivity of hydrogels decreases with the water contents[38,39]. Notably, hydrogels with the equilibrium water contents below 25-30 w/w% exhibit a substantial decrease in the ionic conductivity [40], potentially due to poor percolation between the ion-rich hydrated regions within the hydrogel [41]. Hence, we choose the low water contents hydrophilic polyurethane as an insulating hydrogel for the hydrogel bioelectronic interface. Owing to its low equilibrium water contents around 27 w/w%, the insulating hydrogel exhibits electrical insulating behavior in physiological environments (e.g., PBS) comparable to that of commonly used device encapsulating materials such as PDMS (Fig. S13). Furthermore, the low water contents hydrophilic polyurethane can be dissolved in organic solvent at a high concentration to yield a highly viscous printable ink (Fig. 11b), readily applicable to multi-material printing processes.

**a**

| Conductive hydrogel | Conductivity (S m⁻¹) | Stretchability (%) | Toughness (J m⁻²) | Strength (kPa) | Water contents (%) | Printability | Reference |
|---|---|---|---|---|---|---|---|
| Ionic hydrogels (high concentration salts) | 1-10 | 300-500 | 10-100 | N/R | > 90 | Yes | (*1*) |
| Au nanowire hydrogel | 1-10 | < 100 | N/R | N/R | > 90 | No | (*2*) |
| CNT/graphene hydrogel | 1-10 | < 100 | N/R | N/R | > 90 | No | (*3*) |
| PANi hydrogel (IPN with PVA) | 5-9 | 140 | N/R | 120 | 80-90 | No | (*4*) |
| PEDOT:PSS hydrogel (IPN with PAA) | 23 | 300 | N/R | 50-250 | 50-80 | No | (*5*) |
| PEDOT:PSS hydrogel (IPN with PAAm) | 0.2 | 100 | N/R | 1,200 | 85-90 | No | (*6*) |
| PEDOT:PSS hydrogel (IPN with PVA) | 1.8 | 400 | N/R | 700 | 80-90 | Yes | (*7*) |
| PEDOT:PSS hydrogel (glycerol-based) | 200 | 500 (Plastic deformation) | N/R | N/R | < 10 | No | (*8*) |
| PEDOT:PSS hydrogel (ionic liquid-based) | 4,740 (in DI water) | 30 | N/R | 6 | 82 | No | (*9*) |
| Pure PEDOT:PSS hydrogel (100 w/w% PEDOT:PSS) | 2,000 (in PBS) | 35 | 280 | 500 | 80 | Yes | (*10*) |
| **BC-CPH (25 w/w% PEDOT:PSS)** | **1,100 (in PBS)** | **400** | **3,300** | **900** | **76** | **Yes** | **This work** |

N/R: Not reported

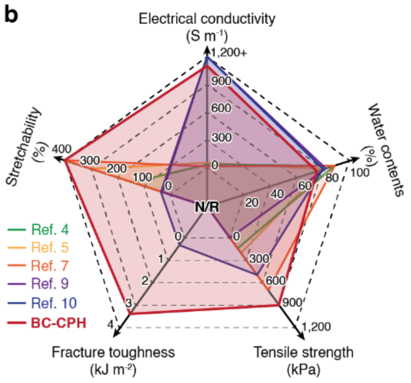

**Fig. S1. Comparison between the BC-CPH and other electrically conductive hydrogels. a** and **b**, Comparison table (**a**) and radar chart (**b**) for various properties of the BC-CPH and the previously reported electrically conductive hydrogels [3,4,6,14-16,18-20,42]. PVA, poly(vinyl alcohol); PAA, poly(acrylic acid); PAAm, polyacrylamide; PANi, polyaniline; PEDOT:PSS, poly(3,4-ethylenedioxythiophene):polystyrene sulfonate.

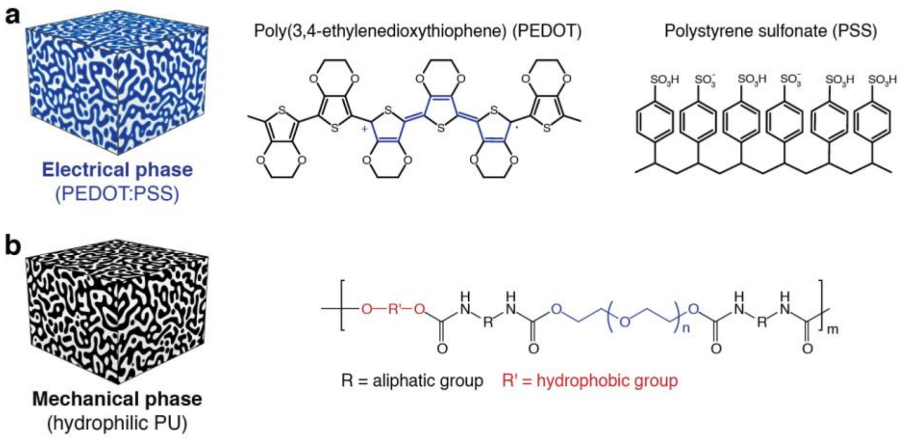


**Fig. S2. Electrical and mechanical phases of the BC-CPH. a**, Electrical phase of the BC-CPH based on poly(3,4-ethylenedioxythiophene):polystyrene sulfonate (PEDOT:PSS). **b**, Mechanical phase of the BC-CPH based on hydrophilic polyurethane (PU).

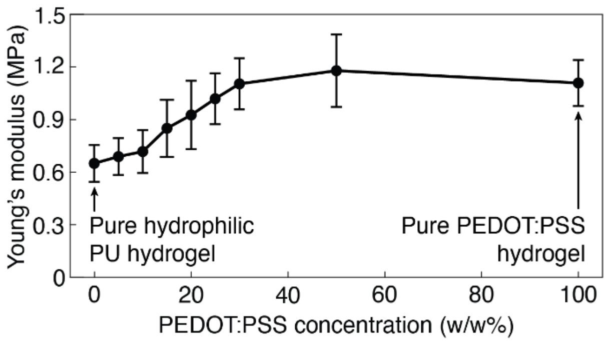


**Fig. S3. Young's moduli of the BC-CPH with varying PEDOT:PSS concentrations.**

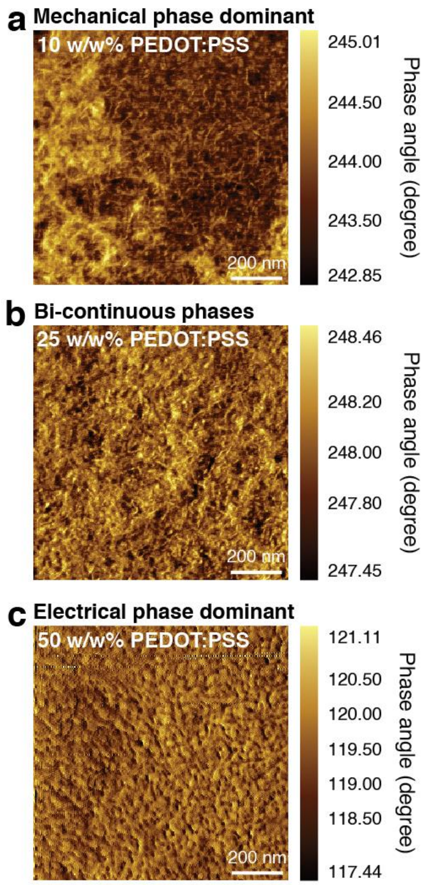


**Fig. S4. AFM phase images of the BC-CPH with varying PEDOT:PSS concentrations. a** to **c**, AFM phase images of the BC-CPH with 10 w/w% (**a**), 25 w/w% (**b**), and 50 w/w% (**c**) PEDOT:PSS concentration.

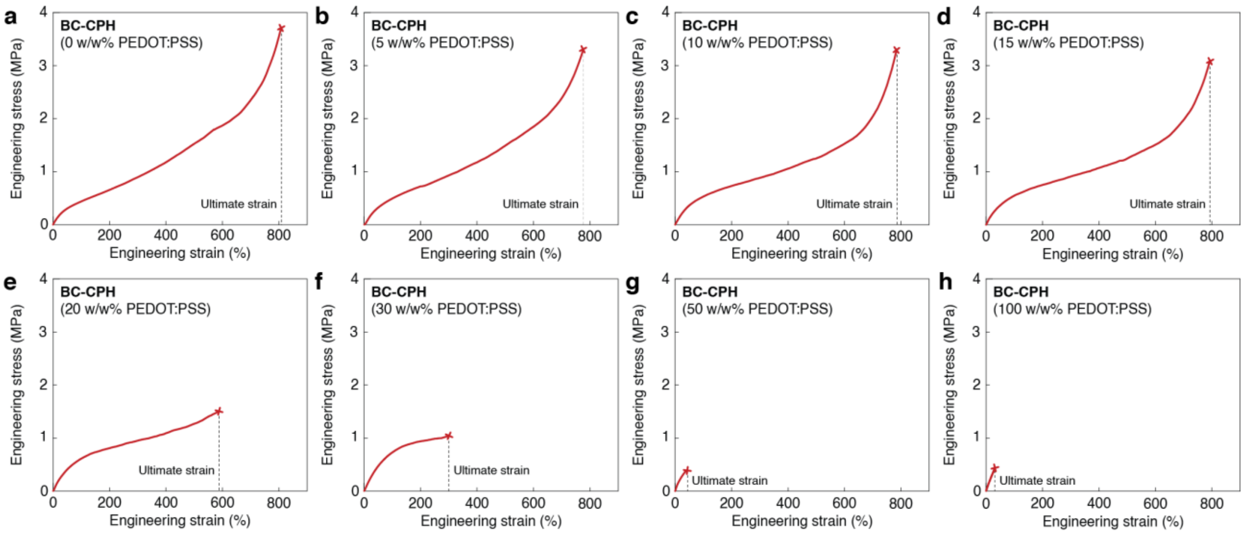


**Fig. S5. Mechanical properties of the BC-CPH with varying PEDOT:PSS concentrations. a** to **b**, Engineering stress vs. engineering strain curves for the BC-CPH with 0 w/w% (**a**), 5 w/w% (**b**), 10 w/w% (**c**), 15 w/w% (**d**), 20 w/w% (**e**), 30 w/w% (**f**), 50 w/w% (**g**), and 100 w/w% (**h**) PEDOT:PSS concentration.

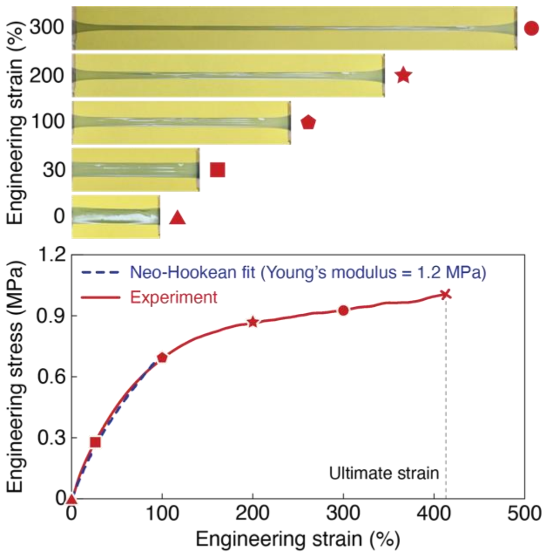


**Fig. S6. High stretchability of the BC-CPH.** Images (top) and plot for engineering stress vs. engineering strain (bottom) for the BC-CPH. Symbols (▲,■,⬟,★,●) on the plot correspond to the images at each strain. The BC-CPH with 25 w/w% PEDOT:PSS is used.

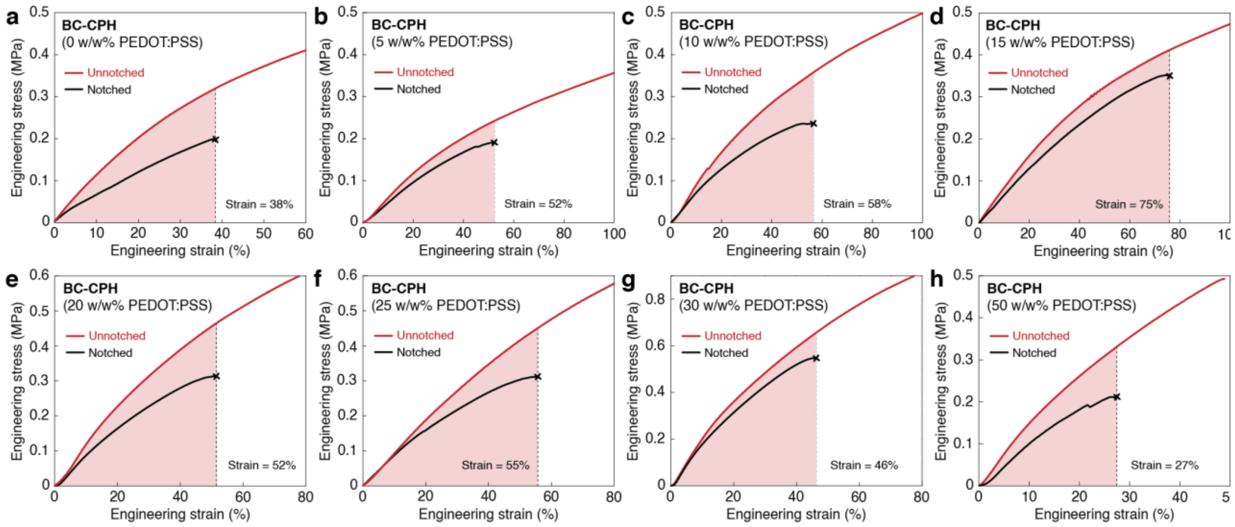


**Fig. S7. Fracture toughness of the BC-CPH with varying PEDOT:PSS concentrations. a** to **h**, Engineering stress vs. engineering strain curves for the unnotched and notched BC-CPH with 0 w/w% (**a**), 5 w/w% (**b**), 10 w/w% (**c**), 15 w/w% (**d**), 20 w/w% (**e**), 25 w/w% (**f**), 30 w/w% (**g**), and 50 w/w% (**h**) PEDOT:PSS concentration.

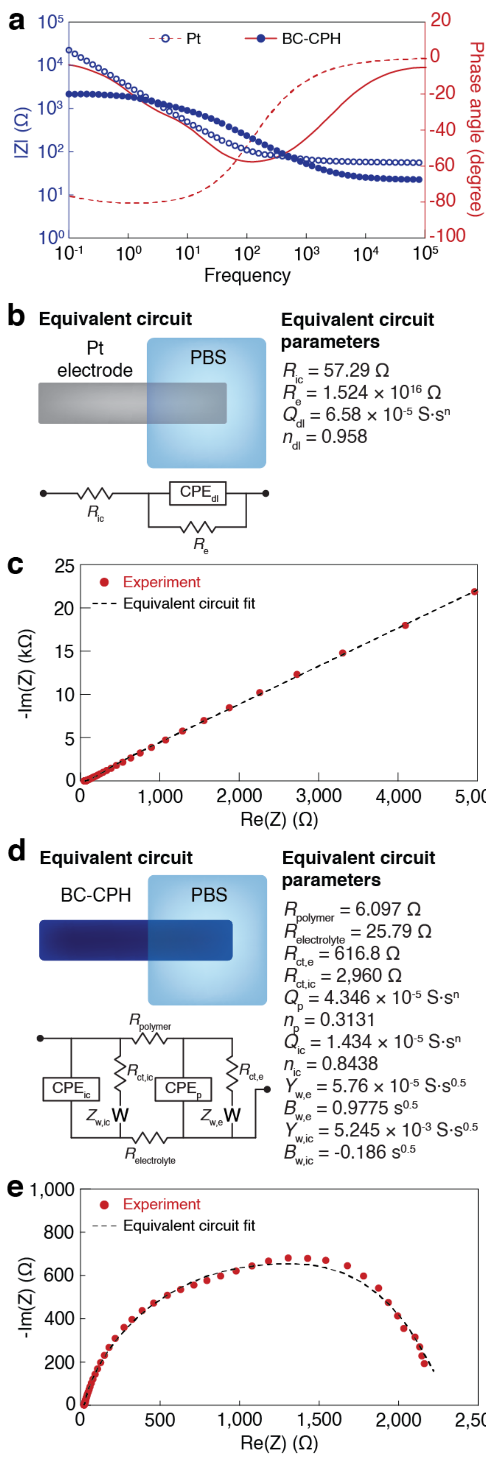


**Fig. S8. Electrical properties of Pt electrode and the BC-CPH. a**, Impedance (circle symbols, left axis) and phase angle (lines, right axis) vs. frequency plots for a Pt electrode and the BC-CPH. **b** and **c**, Equivalent circuit and parameters (**b**) and Nyquist plot (**c**) for a Pt electrode. $R_{ic}$ represents the electronic resistance of the interconnect, $R_e$ represents the electronic resistance of the Pt electrode, and $CPE_{dl}$ represents the double-layer capacitive phase element (CPE) of the Pt electrode. **d** and **e**, Equivalent circuit and parameters (**d**) and Nyquist plot (**e**) for the BC-CPH. $R_{polymer}$ represents the electronic resistance of the BC-CPH, $R_{electrolyte}$ represents the ionic resistance of the electrolyte (PBS), $R_{ct,ic}$ represents the interconnect reaction resistance, $R_{ct,e}$ represents the electrode reaction resistance, $CPE_{ic}$ represents the double-layer CPE of the interconnect, $CPE_{p}$ represents the double-layer CPE of the BC-CPH, $Z_{w,ic}$ represents the interconnect Warburg element, and $Z_{w,e}$ represents the electrode Warburg element. CPE is used to account inhomogeneous or imperfect capacitance and are represented by the parameters $Q$ and $n$ where $Q$ represents the peudocapacitance value and $n$ represents the deviation from ideal capacitive behavior. The true capacitance $C$ can be calculated from these parameters by using the relationship $\boldsymbol{C = Q\omega_{\mathrm{max}}{}^{n-1}}$ where $\boldsymbol{\omega_{\mathrm{max}}}$ is the frequency at which the imaginary component reaches a maximum [43]. Warburg element is used to account diffusion circuit element and represented by the parameters $Y$ and $B$ where $Y$ represents the admittance value and $B$ represents the time constant of the diffusion circuit element. The impedance $Z$ can be calculated from these parameters by using the relationship $\boldsymbol{Z(\omega) = \left(Y\sqrt{j\omega}\right)^{-1} \tanh\left(B\sqrt{j\omega}\right)}$ where $\boldsymbol{\omega}$ is the frequency[44].

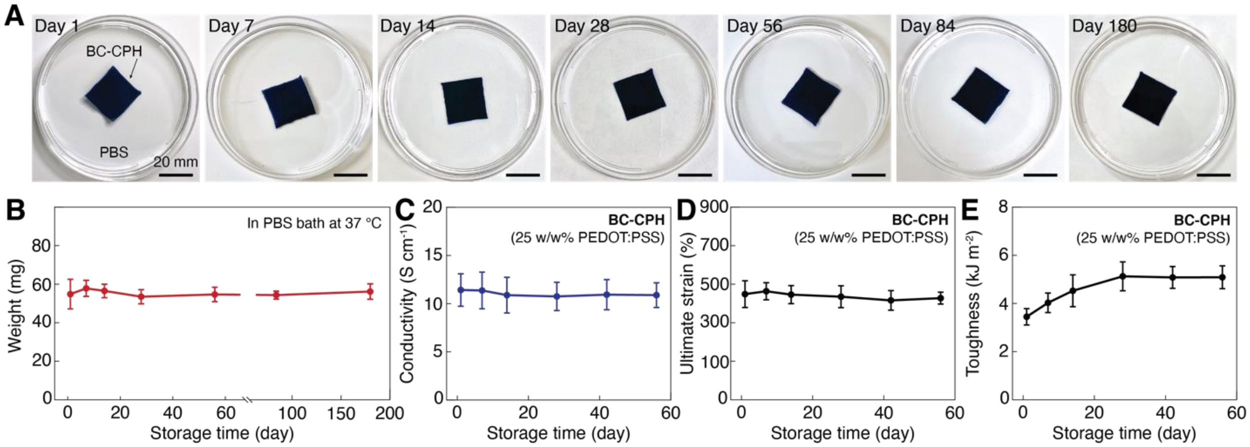


**Fig. S9. Long-term stability of the BC-CPH in physiological environment. a** and **b**, Photographs (**a**) and weight (**b**) of the BC-CPH stored in PBS at 37 °C for 1, 7, 14, 28, 56, 84, and 180 days. **c** to **e**, Electrical conductivity (**c**), ultimate strain (**d**), and fracture toughness (**e**) of the BC-CPH stored in PBS at 37 °C. Error bars indicate SD; $N = 4$.

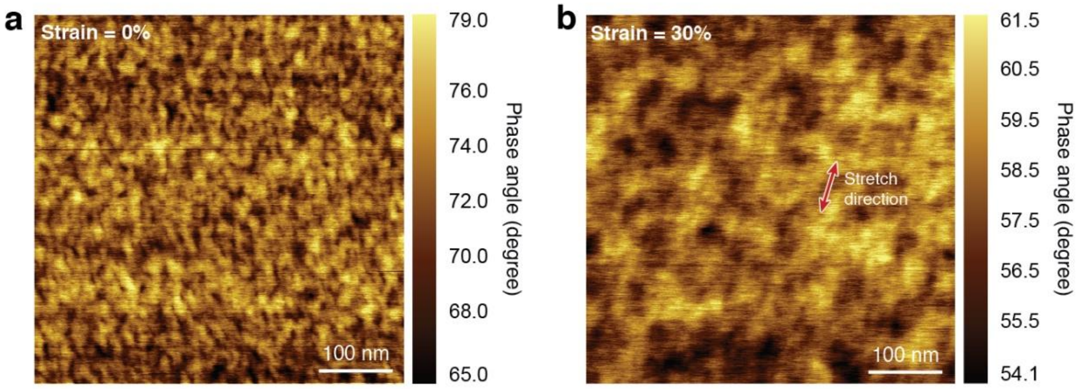


**Fig. S10. Electrical property of the BC-CPH under deformation. a** and **b**, AFM phase images of the BC-CPH at engineering strain of 0% (**a**) and 30% (**b**). Arrow indicates the strain direction. The BC-CPH with 25 w/w% PEDOT:PSS is used.

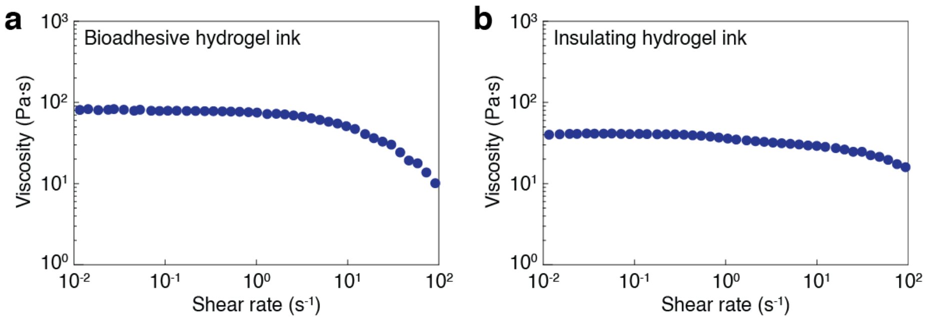


**Fig. S11. Rheological property of the bioadhesive and insulating hydrogel inks. a** and **b**, Viscosity vs. shear rate plots for the bioadhesive (**a**) and insulating (**b**) hydrogel inks.

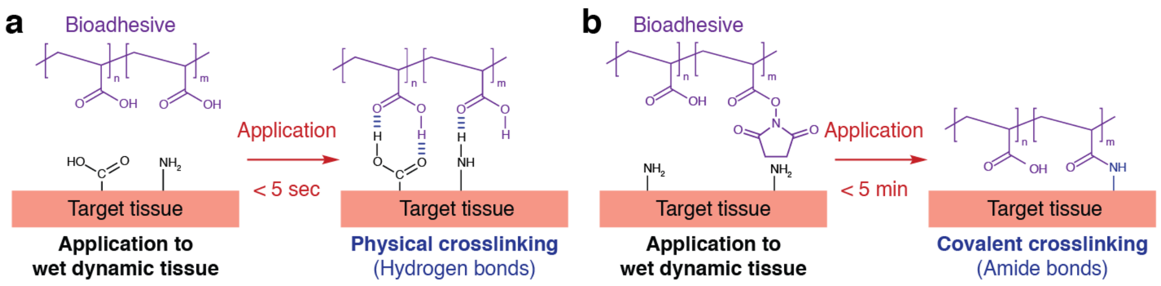


**Fig. S12. Wet adhesion chemistry of the bioadhesive. a**, Schematic illustrations for physical crosslinking between the bioadhesive and the target tissue surface by hydrogen bonds. **b**, Schematic illustrations for covalent crosslinking between the bioadhesive and the target tissue surface by amide bonds.

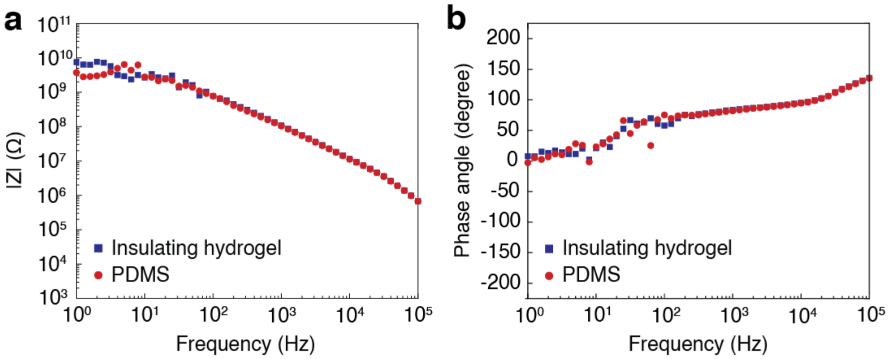


**Fig. S13. Electrochemical property of the insulating hydrogel. a** and **b**, Impedance (**a**) and phase angle (**b**) vs. frequency plots for a PDMS and the insulating hydrogel. The samples with 30 μm in thickness were used for both materials.

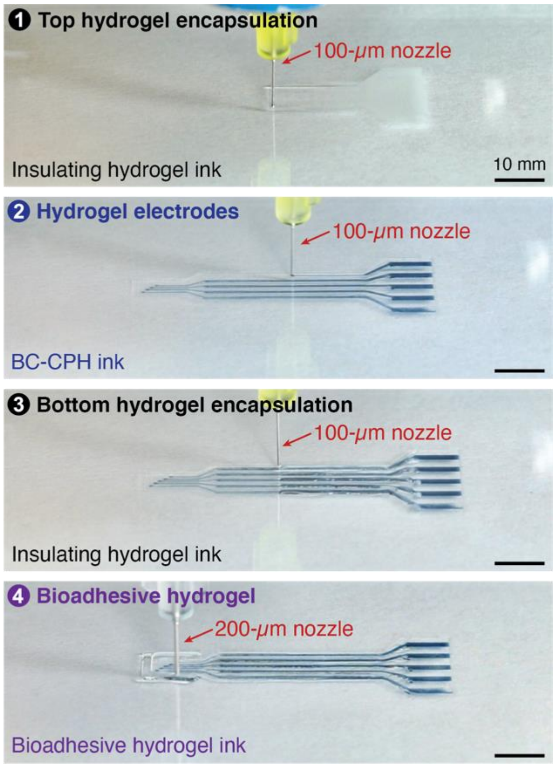


**Fig. S14. Representative printing process of the hydrogel bioelectronic interface.**

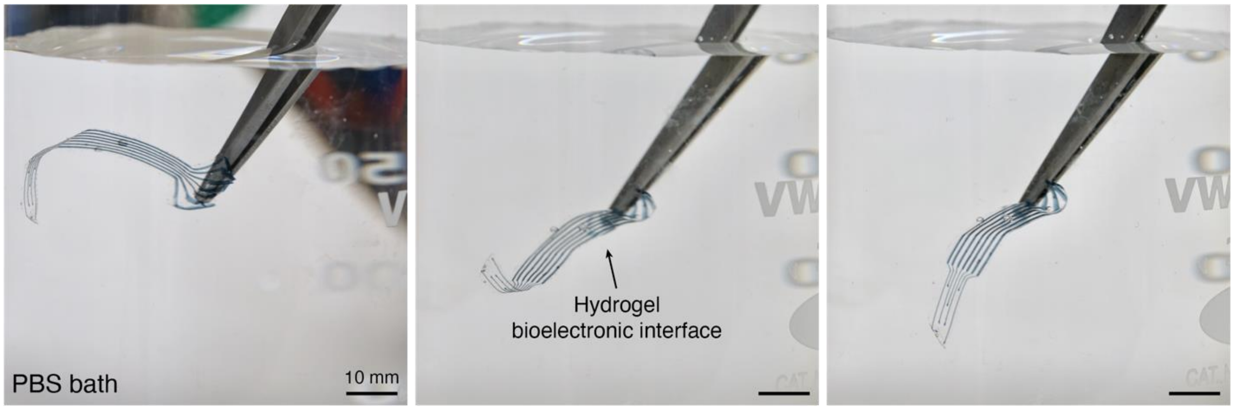


**Fig. S15. Hydrogel bioelectronic interface in PBS bath.**

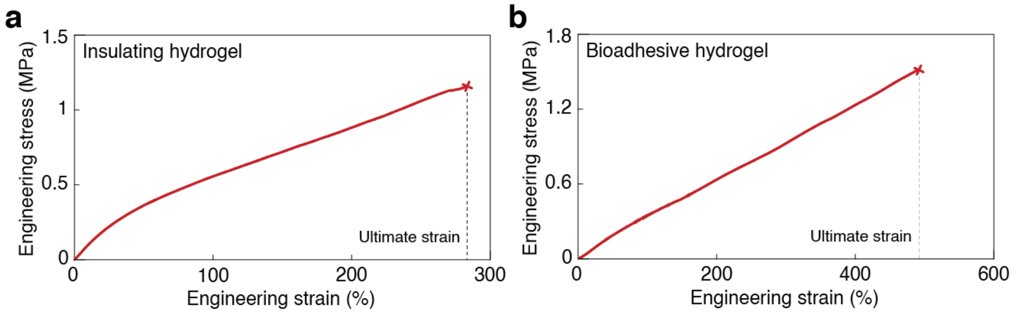


**Fig. S16. Mechanical property of the insulating and bioadhesive hydrogels. a** and **b**, Engineering stress vs. engineering strain plots for the insulating (**a**) and bioadhesive (**b**) hydrogels.

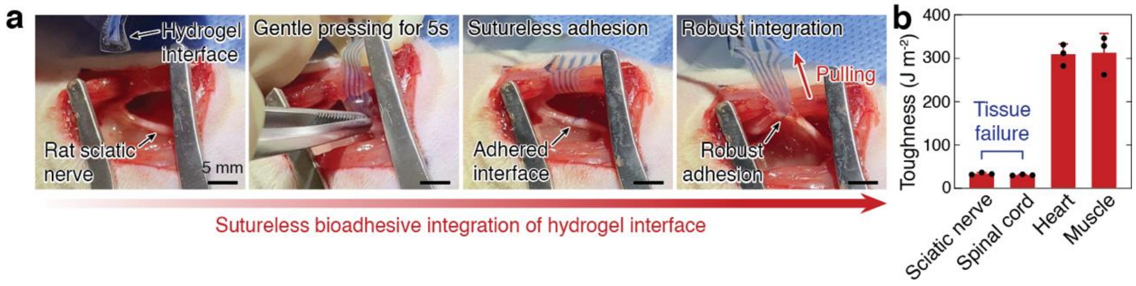


**Fig. S17. Rapid sutureless integration to wet tissues. a**, Snapshots of sutureless bioadhesive integration of the hydrogel bioelectronic interface to a rat sciatic nerve. **b**, Interfacial toughness of the bioadhesive hydrogel adhered to various rat tissues. Note that tissues underwent cohesive failure for sciatic nerve and spinal cord. Error bars indicate SD; $N$ = 3.

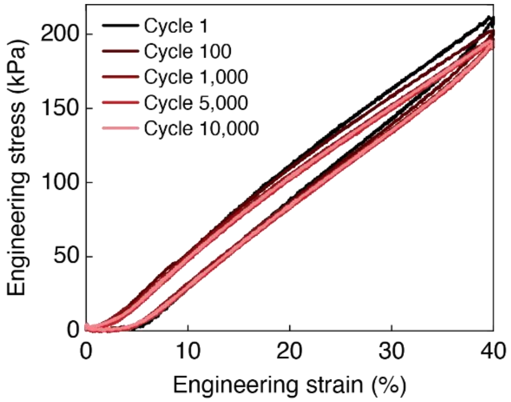


**Fig. S18. Cyclic tensile tests of the hydrogel bioelectronic interface.** Engineering stress vs. engineering strain plots for the hydrogel bioelectronic interface at different cycle numbers.

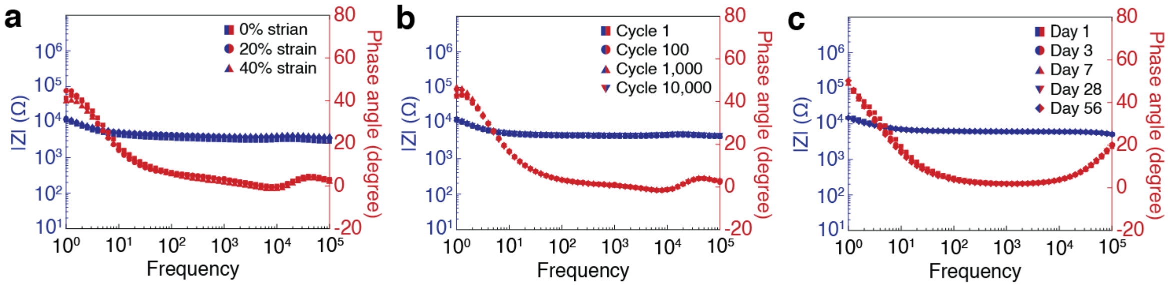


**Fig. S19. Electrochemical stability of the hydrogel bioelectronic interface. a** to **c**, Impedance (blue symbols, left axis) and phase angle (red symbols, right axis) vs. frequency plots for one electrode channel in the hydrogel bioelectronic interface under varying tensile strain (**a**), tensile cycle (**b**), and storage time in a PBS bath at 37 °C (**c**).

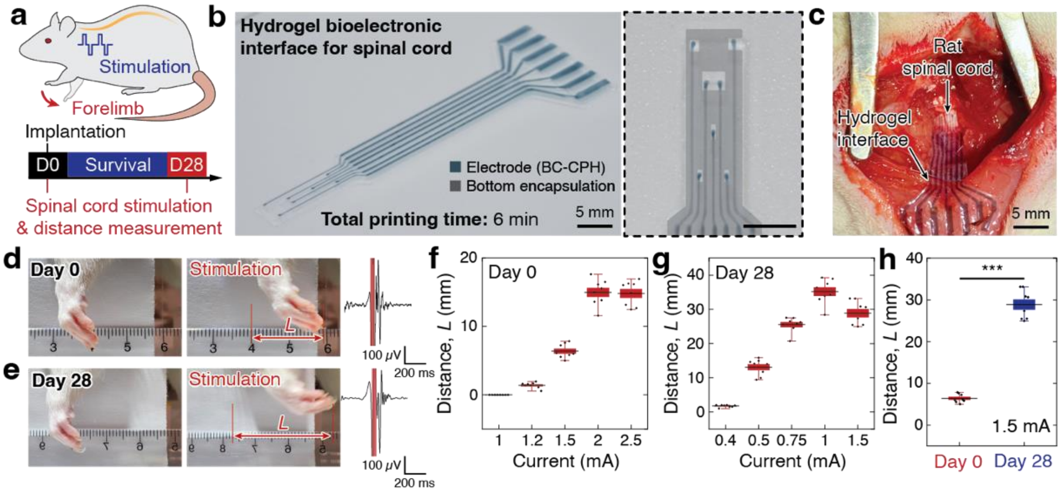


**Fig. S20. Rat spinal cord stimulation by the hydrogel bioelectronic interface. a**, Schematic illustration for rat spinal cord electrophysiological stimulation by the hydrogel bioelectronic interface. **b**, Images of the printed hydrogel bioelectronic interface for spinal cord in the overall view (left) and the magnified view of electrodes (right). Different materials are marked with color overlays in the magnified view. **c**, Images of the implanted hydrogel bioelectronic interface on rat spinal cord. **d** and **e**, Images of rat forelimb before (left) and after (middle) electrophysiological stimulation of the spinal cord by the hydrogel bioelectronic interface with corresponding EMG recordings (right) on day 0 (**d**) and day 28 (**e**) post-implantation. The red-shaded regions in the EMG recordings indicate the stimulation pulses. **f** and **g**, Rat forelimb movement distance upon spinal cord stimulations by the hydrogel bioelectronic interface at varying stimulation currents on day 0 (**f**) and day 28 (**g**) post-implantation. **h**, Comparison of the rat forelimb movement distance on day 0 and day 28 post-implantation with stimulation current of 1.5 mA. In box plots, center lines represent mean, box limits delineate standard error (SE), and whiskers reflect 5th and 95th percentile; $N = 8$. Statistical significance is determined by two-sided Student t-test; *** $P \leq 0.001$.

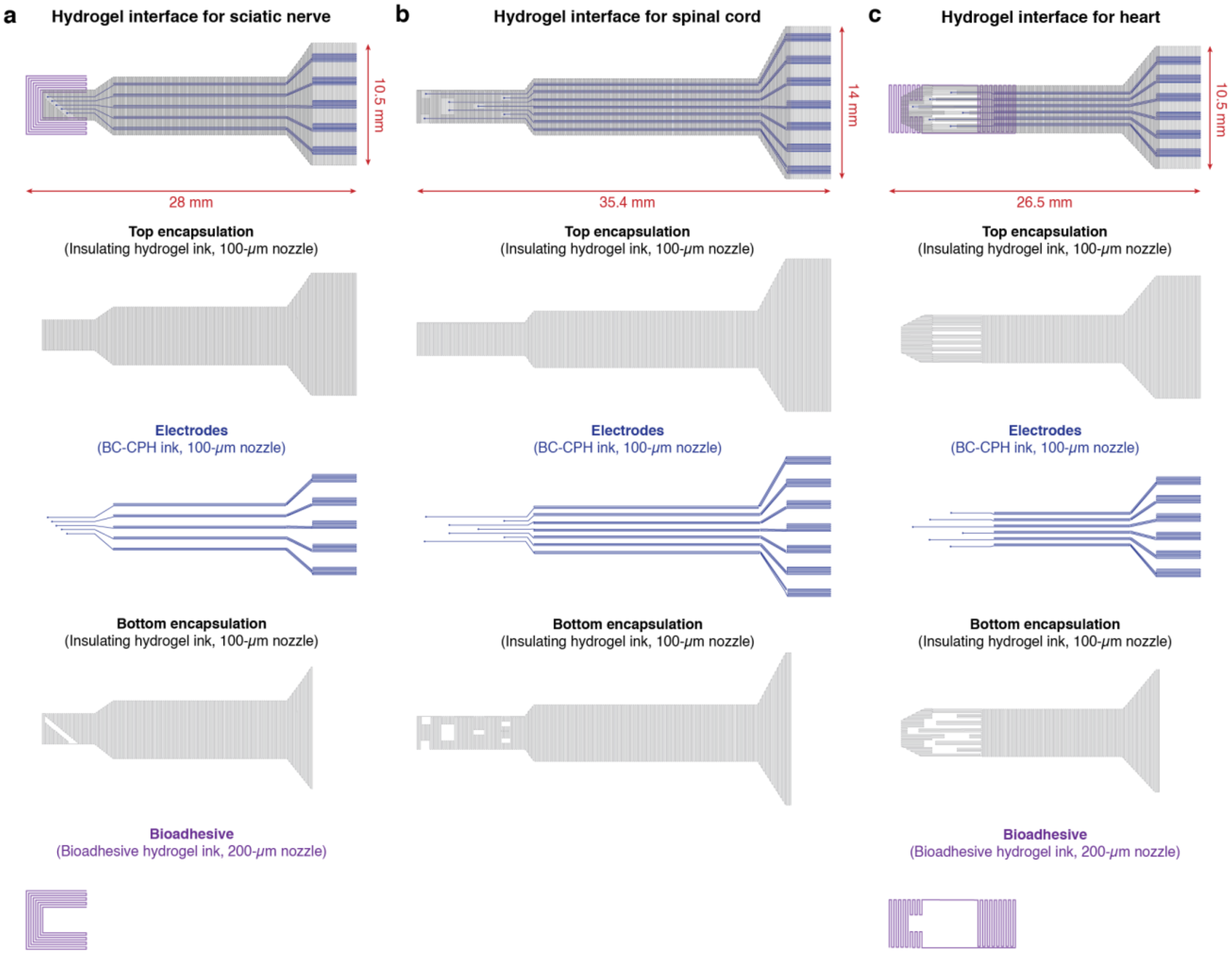


**Fig. S21. Designs for the hydrogel bioelectronic interface for various target tissues. a** to **c**, Overall designs and printing paths for the hydrogel bioelectronic interfaces for sciatic nerve (**a**), spinal cord (**b**), and heart (**c**).

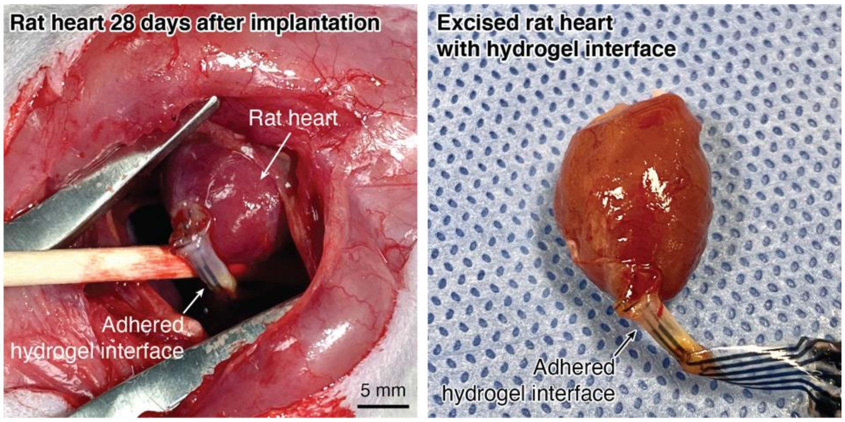


**Fig. S22.** ***In vivo*** **stability of bioadhesive integration of the hydrogel bioelectronic interface.** Images of the hydrogel bioelectronic interface adhered on a rat heart in *in vivo* (left) and in the excised heart (right) on day 28 post-implantation.

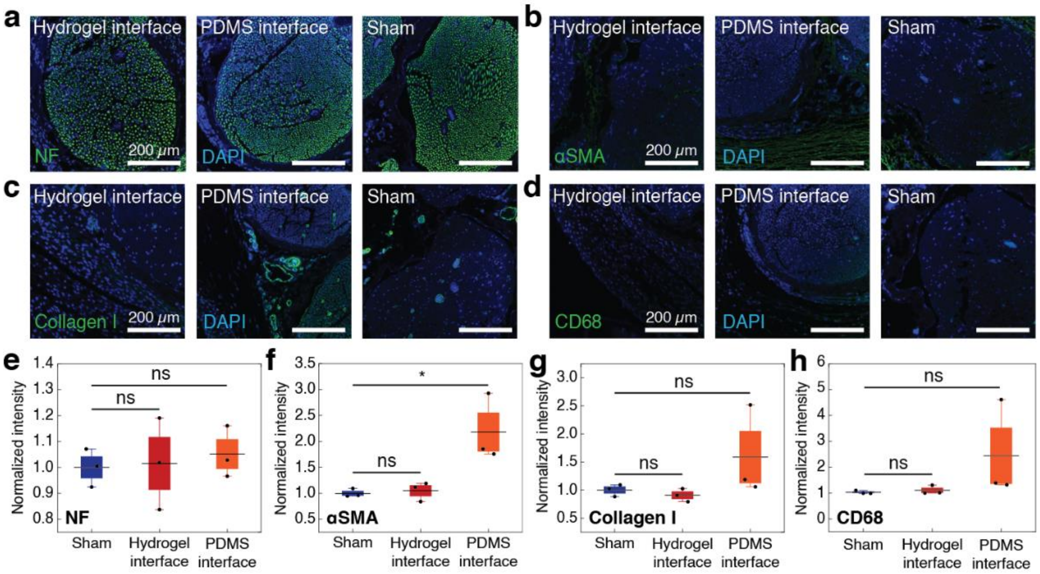


**Fig. S23. Immunofluorescence analysis of rat sciatic nerve on day 7 post-implantation. a** to **d**, Representative immunofluorescence images of rat sciatic nerve on day 7 post-implantation of the hydrogel bioelectronic interface, PDMS interface, and sham group (no device implantation). Cell nuclei are stained with DAPI (blue). Green fluorescence corresponds to the expression of neurofilament (NF, **a**), fibroblasts (αSMA, **b**), collagen (collagen I, **c**), and macrophages (CD68, **d**), respectively. **e** to **h**, Normalized fluorescence intensity plots for the expression of NF (**e**), αSMA (**f**), Collagen I (**g**), and CD68 (**h**) in different groups. In box plots, center lines represent mean, box limits delineate standard error (SE), and whiskers reflect 5th and 95th percentile; $N$ = 3. Statistical significance is determined by two-sided Student t-test; ns, not significant; * $P \leq 0.05$.

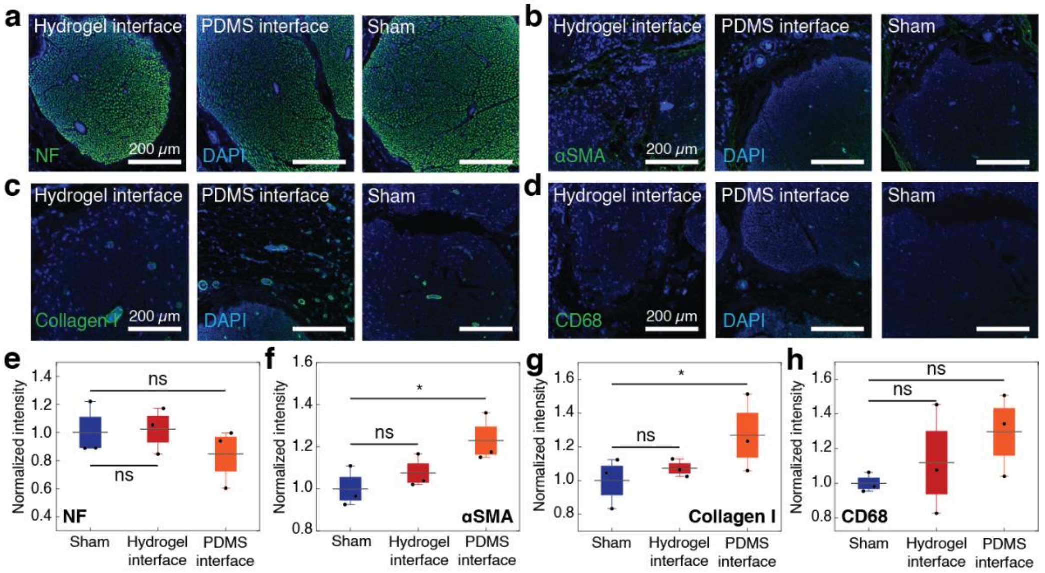


**Fig. S24. Immunofluorescence analysis of rat sciatic nerve on day 56 post-implantation. a** to **d**, Representative immunofluorescence images of rat sciatic nerve on day 56 post-implantation of the hydrogel bioelectronic interface, PDMS interface, and sham group (no device implantation). Cell nuclei are stained with DAPI (blue). Green fluorescence corresponds to the expression of neurofilament (NF, **a**), fibroblasts (αSMA, **b**), collagen (collagen I, **c**), and macrophages (CD68, **d**), respectively. **e** to **h**, Normalized fluorescence intensity plots for the expression of NF (**e**), αSMA (**f**), Collagen I (**g**), and CD68 (**h**) in different groups. In box plots, center lines represent mean, box limits delineate standard error (SE), and whiskers reflect 5th and 95th percentile; $N = 3$. Statistical significance is determined by two-sided Student t-test; ns, not significant; * $P \leq 0.05$.

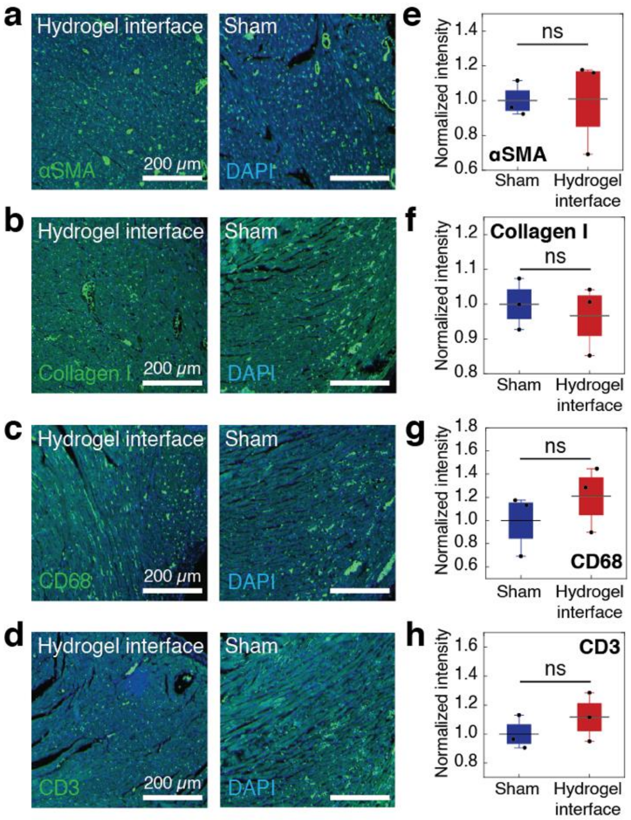


**Fig. S25. Immunofluorescence analysis of rat heart. a** to **d**, Representative immunofluorescence images of rat heart on day 28 post-implantation of the hydrogel bioelectronic interface and sham group (no device implantation). Cell nuclei are stained with DAPI (blue). Green fluorescence corresponds to the expression of fibroblasts (αSMA, **a**), collagen (collagen I, **b**), macrophages (CD68, **c**), and T cells (CD3, **d**), respectively. **e** to **h**, Normalized fluorescence intensity plots for the expression of αSMA (**e**), Collagen I (**f**), CD68 (**g**), and CD3 (**h**) in different groups. In box plots, center lines represent mean, box limits delineate standard error (SE), and whiskers reflect 5th and 95th percentile; $N = 3$. Statistical significance is determined by two-sided Student t-test; ns, not significant; * $P \leq 0.05$.

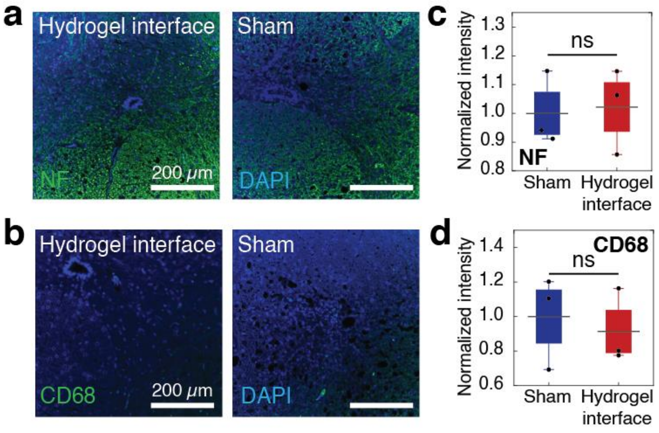


**Fig. S26. Immunofluorescence analysis of rat spinal cord. a** and **b**, Representative immunofluorescence images of rat spinal cord on day 28 post-implantation of the hydrogel bioelectronic interface and sham group (no device implantation). Cell nuclei are stained with DAPI (blue). Green fluorescence corresponds to the expression of neurofilament (NF, **a**) and macrophages (CD68, **b**), respectively. **c** and **d**, Normalized fluorescence intensity plots for the expression of NF (**c**) and CD68 (**d**) in different groups. In box plots, center lines represent mean, box limits delineate standard error (SE), and whiskers reflect 5th and 95th percentile; $N = 3$. Statistical significance is determined by two-sided Student t-test; ns, not significant; * $P \leq 0.05$.

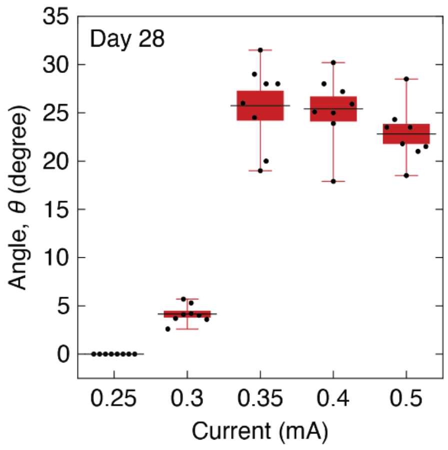


**Fig. S27. Rat sciatic nerve stimulation on day 28 post-implantation.** Rat hindlimb movement angles upon sciatic nerve stimulations by the hydrogel bioelectronic interface at varying stimulation currents. In box plots, center lines represent mean, box limits delineate standard error (SE), and whiskers reflect 5th and 95th percentile; $N = 8$.

**Movie S1.**

Multi-material 3D printing process of a hydrogel bioelectronic interface with the BC-CPH electrodes.

**Movie S2.**

Sutureless bioadhesive integration of a hydrogel bioelectronic interface to a rat sciatic nerve *in vivo*.
